# Genomics of clinal local adaptation in *Pinus sylvestris* under continuous environmental and spatial genetic setting

**DOI:** 10.1101/647412

**Authors:** Jaakko S. Tyrmi, Jaana Vuosku, Juan J. Acosta, Zhen Li, Lieven Sterck, Maria T. Cervera, Outi Savolainen, Tanja Pyhäjärvi

**Affiliations:** Department of Ecology and Genetics, University of Oulu, FI-90014 Oulu, Finland; Biocenter Oulu, University of Oulu, FI-90014 Oulu, Finland; Camcore, Department of Forestry and Environmental Resources, North Carolina State University, Raleigh, NC, USA; Department of Plant Biotechnology and Bioinformatics, Ghent University, Technologiepark 71, 9052 Ghent, Belgium; VIB Center for Plant Systems Biology, Technologiepark 71, 9052 Ghent, Belgium; Centro de Investigación Forestal (CIFOR), Instituto Nacional de Investigaciones Agrarias (INIA), 28040 Madrid, Spain

**Keywords:** adaptation, population genetics – empirical, landscape genetics, gymnosperms

## Abstract

Understanding the consequences of local adaptation at the genomic diversity is a central goal in evolutionary genetics of natural populations. In species with large continuous geographical distributions the phenotypic signal of local adaptation is frequently clear, but the genetic background often remains elusive. We examined the patterns of genetic diversity in *Pinus sylvestris*, a keystone species in many Eurasian ecosystems with a huge distribution range and decades of forestry research showing that it is locally adapted to the vast range of environmental conditions. Making *P. sylvestris* an even more attractive subject of local adaptation study, population structure has been shown to be weak previously and in this study. However, little is known about the molecular genetic basis of adaptation, as the massive size of gymnosperm genomes has prevented large scale genomic surveys. We generated a both geographically and genomically extensive dataset using a targeted sequencing approach. By applying divergence-based and landscape genomics methods we found that several coding loci contribute to local adaptation. We also discovered a very large (ca. 300 Mbp) putative inversion with a signal of local adaptation, which to our knowledge is the first such discovery in conifers. Our results call for more detailed analysis of structural variation in relation to genomic basis of local adaptation, emphasize the lack of large effect loci contributing to local adaptation in the coding regions and thus point out to the need for more attention towards multi-locus analysis of polygenic adaptation.

## Introduction

Populations of species with vast continuous distributions often inhabit very different environments. These populations are often locally adapted, defined as each population having higher fitness than any introduced population at its home site (Kawecki & Ebert, 2004), preferably demonstrated by performing reciprocal transplant experiment (Savolainen, Lascoux, & Merilä, 2013) as has been done with many plant species, such as the *Arabidopsis* genus (e.g. Leinonen, Remington, & Savolainen, 2011, Ågren & Schemske, 2012, Hämälä et al., 2018). Local adaptation can also be inferred from patterns of phenotypic variation or environmental correlation, as has been shown for example, in *Drosophila melanogaster* (Adrion, Hahn, & Cooper, 2015), humans (Fan, Hansen, Lo, & Tishkoff, 2016) and also forest trees (Alberto, Derory, et al., 2013; Savolainen, Pyhäjärvi, & Knürr, 2007).

Local adaptation with a polygenic basis has received more attention lately, because a great deal of adaptive variation is quantitative with multiple underlying loci (Berg & Coop, 2014; Buckler et al., 2009; Rockman, 2012; Yeaman, 2015). A well-known model of polygenic adaptation of a single population in a new environment is presented by the Fisher/Orr model (Fisher, 1918; Orr, 1998) which predicts an exponential distribution of QTL effects (see Nicholas H. Barton & Keightley, 2002). However, local adaptation takes place due to differential selection in different populations in variable environments, possibly connected by gene flow. This kind of selection often results in phenotypic clines (Huxley 1938). Several theoretical predictions for the underlying genetic architecture of clines have been proposed. While differential selection along environmental gradients in continuous populations on single locus governed traits is expected to result in allele frequency clines (e.g. Slatkin, 1973), for polygenic models the expectations are more complex.

Barton (Barton, 1999) has examined a model with polygenic architecture where a subset of loci will have successive, sharp allele frequency clines along the environmental gradient and maintain the phenotypic mean close to the optimum. The underlying loci are, perhaps unrealistically, expected to have similar effect size on the trait. Such sharp allele frequency clines would be seen as F_ST_ outliers, although the majority of loci governing the underlying traits may remain undetected as most alleles are expected to stay near fixation throughout the range in this model.

Latta (Latta, 1998, 2003) and LeCorre and Kremer (Kremer & Le Corre, 2012; Le Corre & Kremer, 2003a, 2012) show that in a high gene flow and strong selection scenario of a polygenic trait, the contribution of covariance between loci becomes more important than between population allele frequency differentiation. Also, in a simulation study Yeaman (2015) has shown that local adaptation is indeed a possible outcome even when only small effect alleles are present given that there is enough standing genetic variation. Most importantly in this model, the contributions of individual loci may be transient making detection of the contributing loci more difficult. Nonetheless, allele frequency clines have been observed in many empirical studies (e.g. (Adrion et al., 2015; Savolainen et al., 2007; Schmidt et al., 2008).

Even with genome-wide datasets of tens or hundreds of thousands of loci sampled across localities, identifying the loci underlying adaptive clinal variation remain a challenge. The majority of methods for uncovering adaptive loci are based on the island model of population structure (Excoffier, Hofer, & Foll, 2009; Foll & Gaggiotti, 2008; Lewontin & Krakauer, 1973; Vitalis, Gautier, Dawson, & Beaumont, 2014) and do not fully utilize the spatial information on the clinal genetic variation. Environmental association analysis (Coop, Witonsky, Di Rienzo, & Pritchard, 2010) and simple regression models can be used to identify clinal trends (e.g. Chen et al., 2012; S T Kujala et al., 2017; Ma, Hall, St. Onge, Jansson, & Ingvarsson, 2010).

It is also important to consider putative effects of recombination across adaptive loci on the genetic architecture of local adaptation. The effect of gene flow is expected to override the effect of weak differential selection on a particular locus. However, physical linkage between multiple small effect alleles makes them behave like a single large effect allele, as described by Yeaman and Whitlock (2011). They show that local adaptation under gene flow may favor genetic architecture where recombination is reduced between loci contributing to local adaptation, which may be caused by physical proximity, transposable element action, translocations or inversions (Kirkpatrick & Barton, 2006). This will result in increased linkage disequilibrium (LD), and thus the examination of unusual LD patterns may be a fruitful approach in discovering the genetic architecture of local adaptation.

The range of *Pinus sylvestris* (Scots pine) spans a huge distribution area in Eurasia from southern Spain to northern Scandinavia and eastern Russia. Its distribution is mostly continuous displaying only limited population structure in the nuclear genome, with the exception of some of the more isolated populations for instance in Spain and Italy (Karhu et al., 1996; Kujala & Savolainen, 2012; Pyhajarvi et al., 2007). However, differentiation within the main range can be seen in mitochondrial haplotype structure, providing information about the recent colonization routes of *P. sylvestris* (Cheddadi et al., 2006; Naydenov, Senneville, Beaulieu, Tremblay, & Bousquet, 2007; T Pyhäjärvi et al., 2007).

In *P. sylvestris* multiple latitudinal phenotypic clines have been repeatedly observed in traits important for abiotic adaptation such as cold tolerance (Aho, 1994; Eiche, 1966; Hurme, Repo, Savolainen, & Pääkkönen, 1997; Hurme, Sillanpää, Repo, Arjas, & Savolainen, 2000) and the timing of growth start and cessation (Beuker, 1994; Karhu et al., 1996; Mikola, 1982). These traits vary latitudinally with environmental conditions, such as temperature, day length, UV radiation intensity and seasonality. Common garden experiments have shown that these traits have a considerable genetic basis suggesting local adaptation (Alberto, Aitken, et al., 2013). When searching for genomic basis of local adaptation, demographic effects may lead to spurious signals if the underlying population structure remans unaccounted for (Hoban et al., 2016). Lack of genome-wide structure, together with highly differentiated phenotypic variation, makes P. sylvestris an ideal species for investigating the genetic basis of local adaptation in large genome. Furthermore, low level of genetic diversity (Kujala & Savolainen, 2012) and lack of any known hybridization with other species should aid in detecting non-equilibrium patterns in the genome. Genetic background of the adaptation remains largely unknown even though some details have been uncovered in previous studies using data from few candidate genes. Only few F_ST_ outliers have been found, but several cases of latitudinal allele frequency clines and variants associated to timing of bud set have been uncovered (Kujala et al., 2017; Kujala & Savolainen, 2012). Similar observations of allele frequency clines have been made in other tree species as well, such as *Populus* (Evans et al., 2014; Ma et al., 2010) and *Picea* (e.g. Chen et al., 2012; Holliday, Ritland, & Aitken, 2010).

Clinal variation, genetic differences across the range and the effect of natural selection in *P. sylvestris* are obvious at the phenotypic level. In this study we use the first genome-wide dataset of genetic diversity in *P. sylvestris* to answer the following questions regarding the manifestation of the phenotypic patterns at the molecular level of variation: 1) Are there deviating levels of genome-wide genetic diversity or LD patterns indicating the impact of local selection? 2) Have dispersal patterns (wind pollination) and the continuous distribution, common in tree species, resulted in spatially continuous isolation-by-distance pattern across the genome and distribution range? 3) Do we see genomic signatures of local adaptation in the form of allele frequency clines, F_ST_ outliers or differentiation of structural variation in *P. sylvestris*?

## Materials and methods

### Plant material and genotyping

Seeds from 109 *P. sylvestris* samples from 12 populations spanning 31 degrees of latitude were used in generating the dataset for this study (Figure 1, Table 1). The main sampling area included two latitudinal gradients, one from northern Finland to Poland and another north-south gradient in western Russia. A total of 120 samples were initially genotyped, of which one was later removed due to sampling the same tree twice and additional 10 were removed due to low sequencing coverage. Haploid genomic DNA was extracted from megagametophyte tissue by using E.Z.N.A.^®^ SP Plant DNA kit (Omega Biotek). DNA was fragmented to an average length of 200 nucleotides with Bioruptor® ultrasonicator (Diagenode). Libraries were prepared by using NEBNext® DNA Library Prep Master Mix Set for Illumina and NEBNext Multiplex Oligos for Illumina E7600S (New England BioLabs) for multiplex sequencing, multiplexing libraries of four samples. Targeted capture was performed for each pool according to MycroArray MYbaits protocol v.2.3.1.

**Figure 1.**
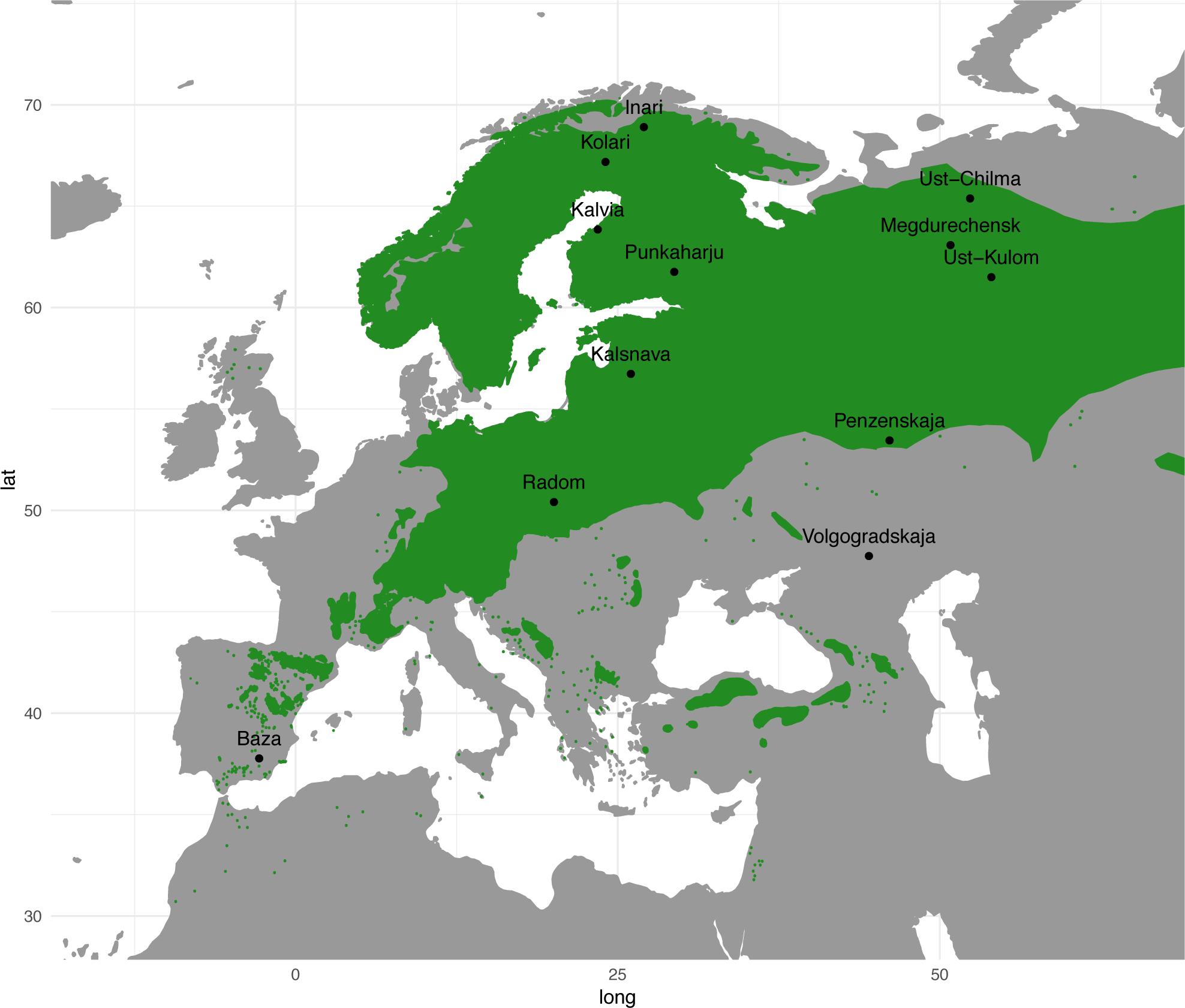
Map of sampling locations with P. sylvestris distribution marked with green color.

**Table 1.**
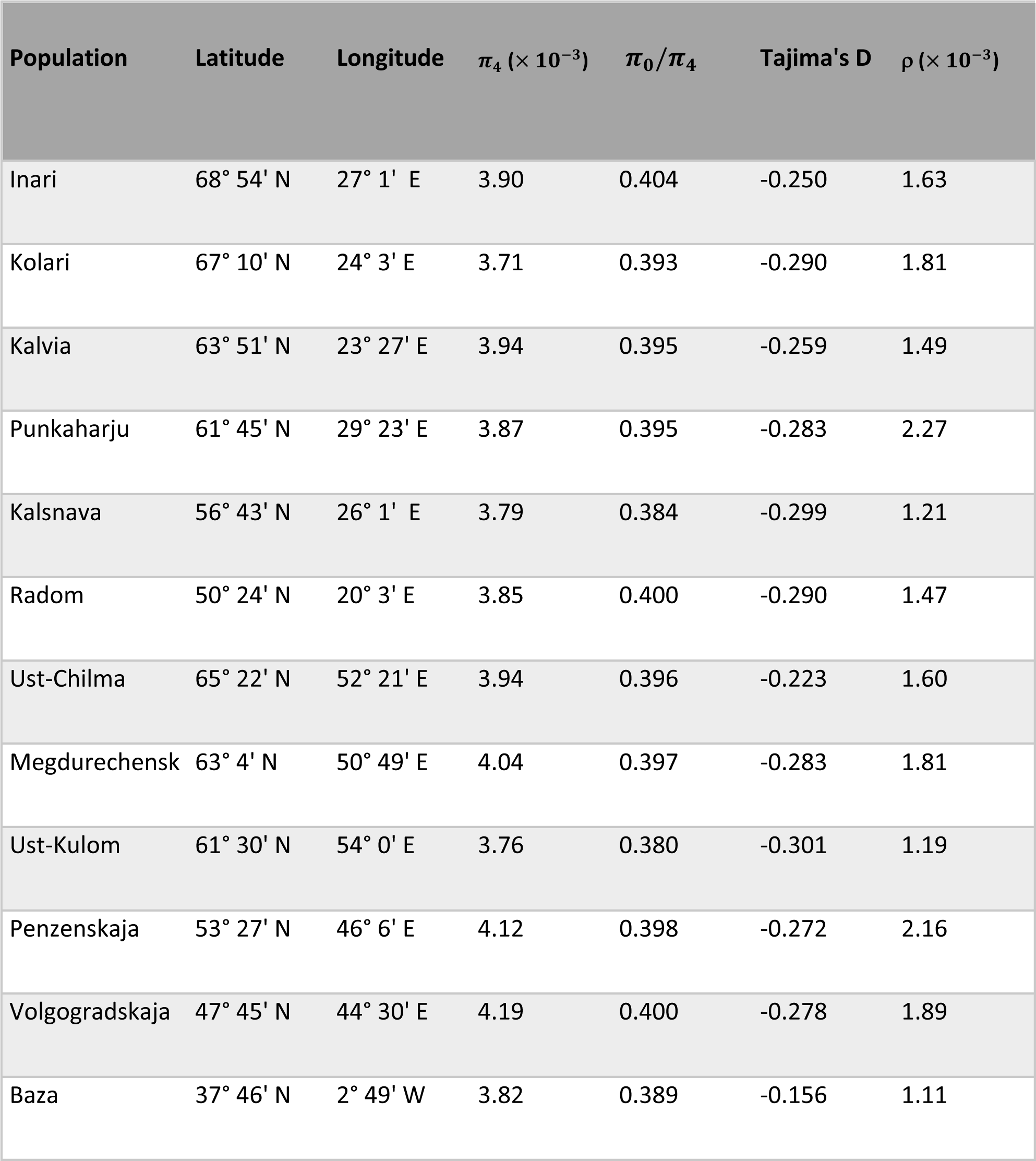
Study populations location and summary statistic information.

### Bait design for targeted sequence capture

The genome size of the *P. sylvestris* is about 23.6 Gbp (Zonneveld, 2012) and there is no reference genome available. As whole genome resequencing approaches are not economically feasible at large scale, baits targeting genic regions together with Illumina paired end sequencing were used to assess allele frequency patterns at genome wide level. Bait design was based on a set of *P. sylvestris* transcripts described previously (Z. Li et al., 2017). Briefly, transcriptomes of *P. sylvestris* were assembled from 454 read data derived from different developmental stages using the Newbler software (v2.8.1). Those were then integrated with public transcriptomes from PlantGDB-assembled Unique Transcripts (based on GeneBank release 187) and a public set of EST assemblies. This initial set contains 121,538 transcripts on which 36,106 open reading frames (ORFs) were predicted by TransDecoder (r20131117). The ORFs were then mapped against the repeat masked *P. taeda* reference genome version 1.01 (Neale et al., 2014) with gmap (Wu & Nacu, 2010) in order to obtain exon sequences. An ORF was omitted if it could be mapped equally well to several locations of the reference suggesting a paralogous sequence. In total 10,330 ORFs encompassing an area of 12,221,835 bp were selected as targets for initial bait design.

MycroArray MYbaits (Ann Arbor, MI) service was used to create an initial set of 100 base long baits with 2x tiling resulting in a total of 176,334 baits. Four pilot experiments including target capture and sequencing were then conducted to determine bait performance. The putative position of each bait in the genome was determined by aligning the bait sequences to the *P. taeda* reference genome v. 1.01 (Neale et al., 2014) with blastn. A well working bait was defined as having a unique high-quality hit to omit possible paralogous sequences, at least 75 out of 100 bases aligning to omit baits on exon-intron boundaries and less than 4 % mismatches to ensure successful alignment. To analyze bait ability to capture target areas 2 x 100 bp paired-end sequencing reads were generated in total for 32 *P. sylvestris* megagametophyte and needle samples with Illumina Hiseq 2500 instrument and 2 x 150 bp paired-end reads with MiSeq instrument. Baits failing to capture any sequence were omitted from the final bait set. After filtering 60,000 high quality bait sequences were selected as the final bait set that was used for examining the 109 samples used in this study.

### Genotype calling workflow

Raw reads from Illumina sequencing were aligned to *P. taeda* reference genome version 1.01 with bowtie2 version 1.1.1 (Langmead & Salzberg, 2012) with parameters --no-mixed --no-discordant --no-unal. The resulting SAM files were modified with Picard toolkit (http://broadinstitute.github.io/picard/) and SAMTools (H. Li et al., 2009) by converting SAM files to BAM format with SamFormatConverter, sorting with SortSam, removing duplicate sequences with MarkDuplicates, defining read groups with AddOrReplaceReadGroups and indexing with SAMTools index. Examination of alignments revealed that the repetitive nature of the *P. sylvestris* genome was causing issues in read alignment and possibly leading to spurious SNP calls. The process of detecting the issue and circumventing incorrect SNP calls is described in supplementary methods. In short, the SNP calling was performed twice using freebayes, first to detect problematic areas identified as heterozygous SNP calls not expected when sequencing haploid DNA, and second time to call SNPs in only in problem-free areas.

The technical quality was evaluated by generating a fastqc quality report for raw reads, samtools flagstat report for alignment success, along with visual inspection of alignments with samtools tview and Integrative Genomics Viewer (Thorvaldsdóttir, Robinson, & Mesirov, 2013). Based on these reports, 10 samples were removed due to low technical quality. Variant calls were filtered with VCFtools (Danecek et al., 2011) to remove sites with quality score below 30 and read depth lower than 5. The entire variant position was removed if it contained non-SNP variants, non-biallelic variants, or had more than 33 % missing data. The final high-quality dataset contained 81,301 SNPs. The genotype calling workflow was parallelized using workflow management software STAPLER (Tyrmi, 2018).

### Diversity and population structure

To estimate the levels of genetic diversity, pairwise nucleotide diversity (Nei & Li, 1979) was calculated with a modified version of python script provided in Garner et al. (2016). The size of available genome used for analysis was 3.8 Mbp. To calculate Tajima’s D and pairwise F_st_the SNP data set was filtered with vcftools --thin parameter to remove variants closer than 10 kbp from each other to reduce correlation between sites due to physical proximity. After filtering, a set of 4,874 SNPs were available.

Tajima’s D estimates (Tajima, 1989) were calculated for the whole dataset and also for each population separately using ∂a∂i (Gutenkunst, Hernandez, Williamson, & Bustamante, 2009). The allele frequency spectrum was generated using ∂a∂i with first down-projecting the sample size to 86 to account for missing data. Hudson’s (1992) pairwise F_ST_ values were then calculated for each population pair by using the equation developed by Bhatia et al. (2013) for a two-population, two locus scenario. An unbiased genome-wide estimate of F_ST_ for each population pair was obtained by calculating the nominator and denominator of equation 10 presented in Bhatia et al. (2013) separately for each site, averaged over all sites after which the division was performed.

Population structure was also examined with principal component analysis (PCA) (McVean, 2009) using the prcomp R package. Sample size was evened between different populations to six as uneven sample sizes may distort the PCA projections. For the analysis singleton variants were removed as recommended for instance by Galinsky et al. (2016).

Population structure was further analyzed by using STRUCTURE software (Pritchard, Stephens, & Donnelly, 2000). As STRUCTURE is computationally demanding the SNP set was stringently filtered to obtain a smaller high-quality data set of 4,197 bi-allelic SNPs. At least 10 kbp distance between variants, minor allele frequency of 0.3 and maximum proportion of missing data per site of 0.2 was used. Burn-in length of 250,000 and run length of 50,000 steps were used. K values from 1 to 10 were tested with three replicate runs for each value of K. The software package Clumpak (Kopelman, Mayzel, Jakobsson, Rosenberg, & Mayrose, 2015) was used to visualize the results and in determining the most likely value of K by using the method of Evanno et al. (2005).

The R-package conStruct (Bradburd, Coop, & Ralph, 2018) was used for spatial analysis of population structure. It allows explicit testing for presence of isolation-by-distance, often found in continuous populations and thus reduces the probability of overestimating the number of potential clusters. Two models, non-spatial which is similar to the model the ADMIXTURE software uses (Alexander, Novembre, & Lange, 2009), and a spatial model which accounts for isolation-by-distance patterns, were tested. For both models K-values from 1 to 5 were examined using 50,000 iterations in each. To test whether the spatial or non-spatial model better explains the genetic variation and to compare results between different K-values the cross-validation pipeline provided with conStruct was used with 50 iterations up to K value of 8 (Figure S1).

The level of LD in the dataset was estimated with the allelic frequencies correlation coefficient *r^2^* (Hill & Robertson, 1968). *r^2^* was calculated between all variants located within the same scaffold over all populations as the populations are nearly panmictic according to the conStruct analysis. Singletons were omitted from this analysis.

To detect loci forming allele frequency clines along the sampled latitudinal gradients, possibly indicating that they are under varying selective pressure along the gradient, a test of generalized linear mixed effect models was fitted for all loci using R package lme4. The first model was created with glmer function for each SNP by setting genotypes as a response variable, population information as a fixed effect and latitude as a random effect. The second model was created similarly but with latitude omitted. The two models were then compared to each other by calculating a p-value with ANOVA to infer whether or not latitude contributes to the model. The Baza population was omitted from all selection scan analyses as it was shown to be the only population clearly differentiated from the others in every analysis of population structure. In addition the method bayenv (Coop et al., 2010) was tested, but the results are not reported because extreme inconsistencies were observed between independent runs that included population covariance structure, indicating a high risk of false positive outliers.

To identify putative loci responsible for local adaptation we used the program pcadapt (Luu, Bazin, & Blum, 2017). It infers population structure with PCA and then identifies putative outliers with respect to how they are related to the population structure, making it well suited for examining datasets containing isolation-by-distance patterns. We used all SNPs with minor allele count over 10 and with Baza population omitted for generating PCA. The number of principal components to be used in the outlier analysis was chosen as two, by first producing a scree plot (Figure S2) with pcadapt and then applying Cattell’s graphical rule.

To further detect potential loci underlying local adaptation, we also used the Bayesian FST-outlier method bayescan (Foll & Gaggiotti, 2008) that is based on identifying locus specific components affecting allele frequencies as a signal of selection. Bayescan was run with default parameters with the exception of setting prior odds for the neutral model to 100 from the default of 10 to account for the large set of SNPs as per the recommendation of bayescan documentation.

Bayescan analysis also revealed the presence of a large haplotype structure in 11 samples with SNPs in complete LD in several scaffolds. To find all scaffolds included in the haplotype structure an r^2^ value was calculated between one of the SNPs contained in the haplotype and all other SNPs in our dataset. Scaffolds containing one or more SNPs with r^2^ value of 1.0 in this comparison were then assumed to be part of the haplotype structure. As it is possible that SNPs are in complete LD only by chance, we examined how likely it is that the whole haplotype structure would be due to chance. This was done by randomly choosing 10,000 sets of scaffolds with each set having similar properties to the ones containing the haplotypes and testing how often similar haplotype structure could be seen. More specifically, each randomized set contained similar number of scaffolds (59) with similar number of SNPs (25 per scaffold) and contained at least one SNP with equally high or higher minor allele frequency (11/109) as the scaffolds containing the haplotype.

Raw Illumina sequences are openly available at NCBI SRA with accession number (TBA). Scripts and relevant output files are accessible at Dryad (TBA) (Tyrmi et al. 2019).

## Results

### Nucleotide diversity

Pairwise synonymous nucleotide diversity averaged over populations was 0.0039 with different populations showing similar diversity (Table 1). *π_N_*/*π_S_* ratio was on average 0.394 and again similar levels can be seen in all populations indicating homogenous levels of negative selection across populations. Tajima’s D value over all populations was −1.29 and it was also negative within every sampled population with Baza population having a higher value compared to others (Table 1). This result is also reflected in the minor allele frequency spectrum calculated for all samples (Figure 2), which shows an excess of rare alleles compared to the standard neutral expectation.

**Figure 2.**
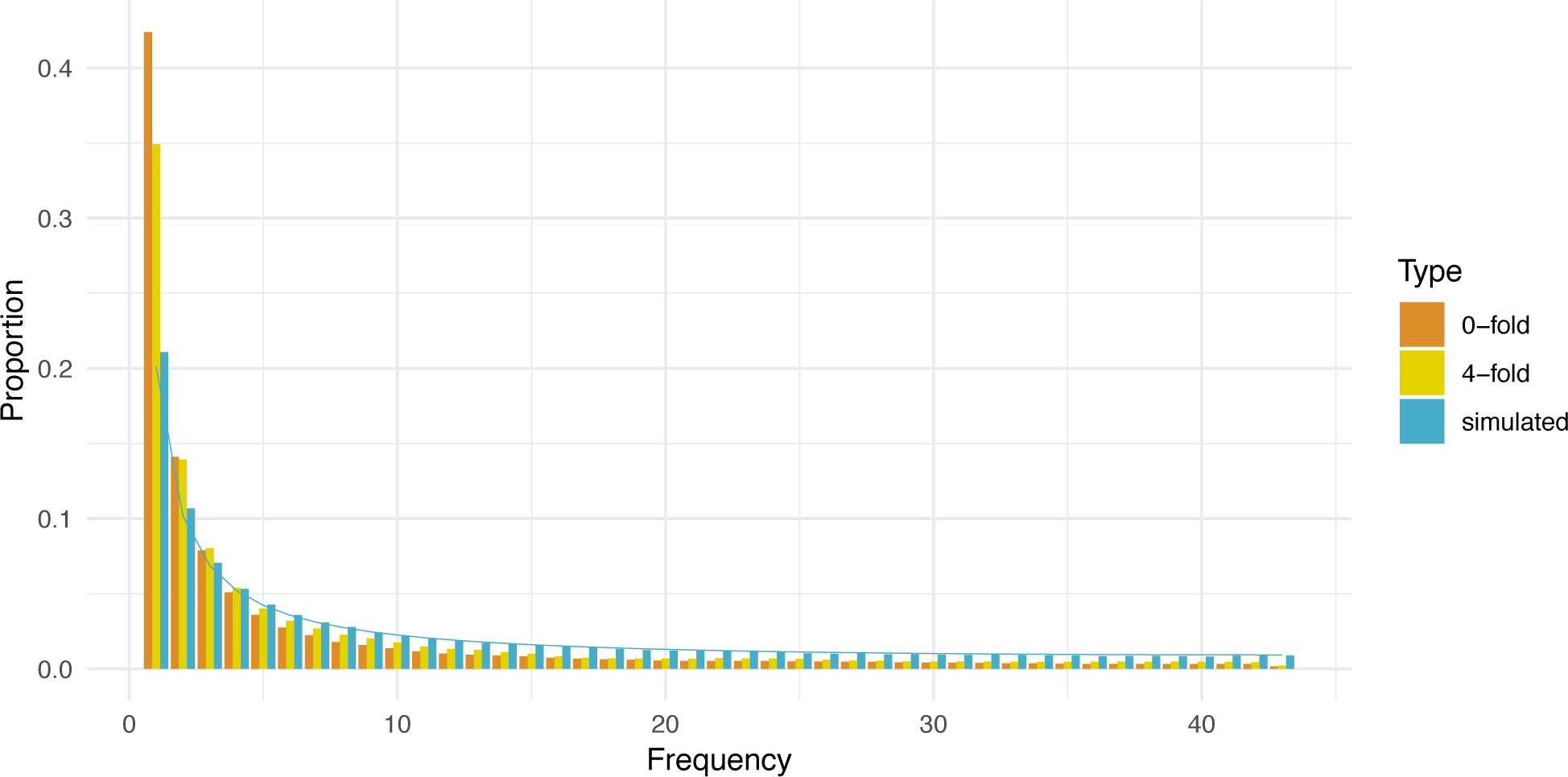
Minor allele frequency spectrum calculated over all samples. Spectrum is projected down to 87 samples to account for missing data

### Assessing population structure

We performed a pairwise F_ST_ (Hudson et al., 1992), STRUCTURE (Pritchard et al., 2000) and PCA (McVean, 2009) analysis to evaluate the genetic relationships between populations. All analysis indicate that the Spanish Baza population is differentiated from other populations and in addition to that, very subtle population structure separates eastern and western samples from each other. In general, pairwise F_ST_ estimates show low level of differentiation between most sample populations (Table 2) with overall F_ST_ of 0.031. Contrasts with Baza are higher (average of 0.079).

**Table 2.** Weighted genome-wide averages of pairwise F_ST_ estimates for all populations. Higher values are marked with darker shades of grey.

| Population | Inari | Kolari | Kalvia | Punkaharju | Kalsnava | Radom | Ust-Chilma | Megdurechensk | Ust-Kulom | Penzenskaja | Volgogradskaja |
| --- | --- | --- | --- | --- | --- | --- | --- | --- | --- | --- | --- |
| Kolari | 0.013 |  |  |  |  |  |  |  |  |  |  |
| Kalvia | -0.001 | 0.016 |  |  |  |  |  |  |  |  |  |
| Punkaharju | 0.000 | 0.017 | 0.005 |  |  |  |  |  |  |  |  |
| Kalsnava | 0.019 | 0.019 | 0.023 | 0.020 |  |  |  |  |  |  |  |
| Radom | 0.002 | 0.021 | 0.005 | 0.007 | 0.015 |  |  |  |  |  |  |
| Ust-Chilma | 0.010 | 0.022 | 0.013 | 0.010 | 0.034 | 0.018 |  |  |  |  |  |
| Megdurechensk | 0.008 | 0.021 | 0.011 | 0.009 | 0.029 | 0.016 | 0.001 |  |  |  |  |
| Ust-Kulom | 0.064 | 0.045 | 0.063 | 0.064 | 0.044 | 0.052 | 0.054 | 0.052 |  |  |  |
| Penzenskaja | 0.009 | 0.023 | 0.011 | 0.013 | 0.033 | 0.015 | 0.008 | 0.009 | 0.060 |  |  |
| Volgogradskaja | 0.000 | 0.017 | 0.005 | 0.005 | 0.023 | 0.007 | 0.010 | 0.006 | 0.063 | 0.000 |  |
| Baza | 0.068 | 0.082 | 0.073 | 0.071 | 0.077 | 0.065 | 0.083 | 0.077 | 0.117 | 0.081 | 0.072 |

The STRUCTURE results (Figure 3) analyzed using Evanno method indicate that the most likely value of K is three. Again, Baza population forms a distinct group compared to the main distribution. However, the rest of the range is divided into two groups, where the other contains all samples of Ust-Kulom population and parts of other, mainly eastern, populations. PCA analysis suggests that the genetic differentiation within and across populations in general is continuous as each principal component explains just small fraction of total variance (Figure 4A), although some trends can be observed. The samples originating from Baza population are separated from the rest by the first principal component. The rest of the range is being clustered more closely together, with a trend separating the eastern and western samples from each other (Figure 4B, 4C).

**Figure 3.**
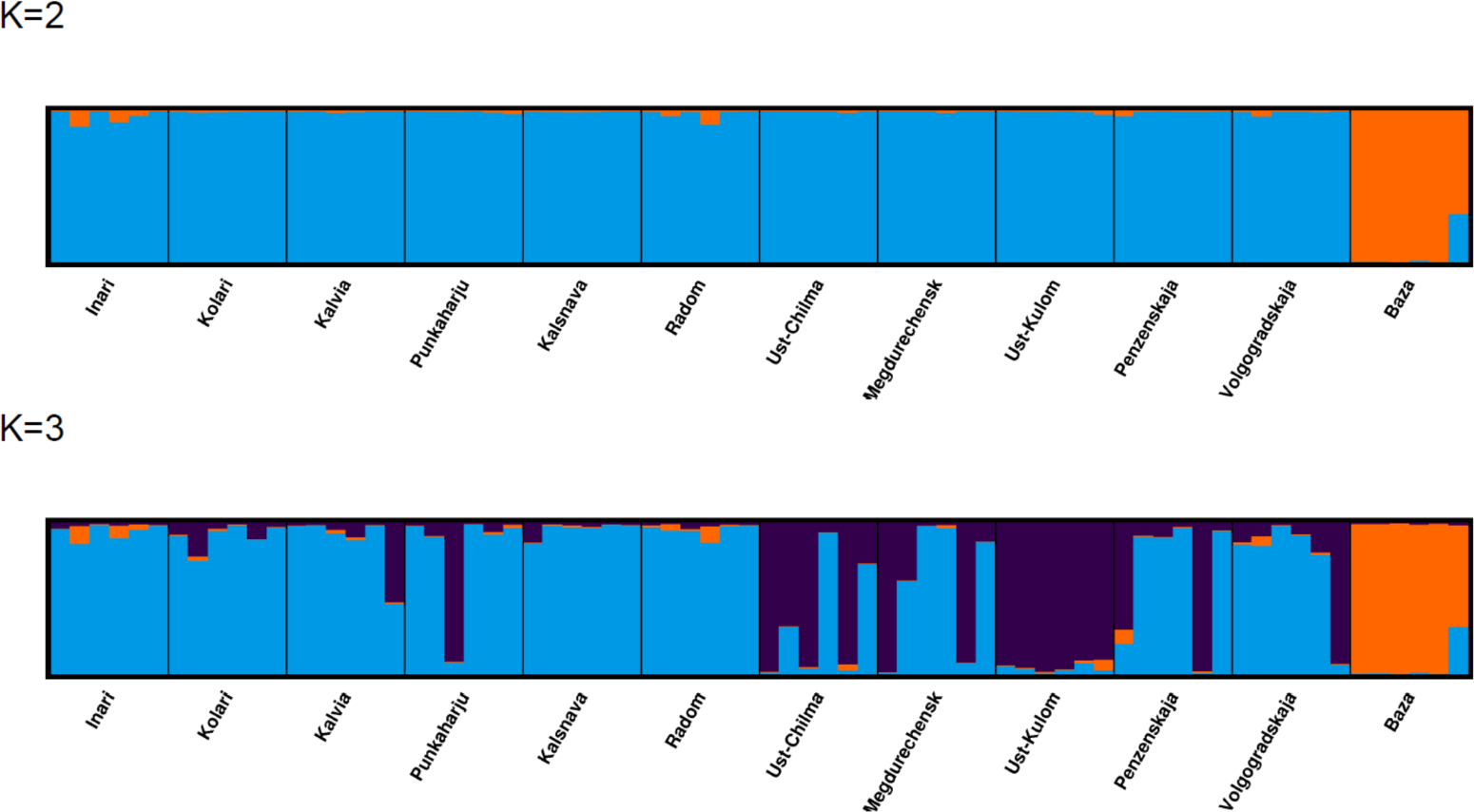
Visualization of STRUCTURE results using K values of 2 and 3.

**Figure 4.**
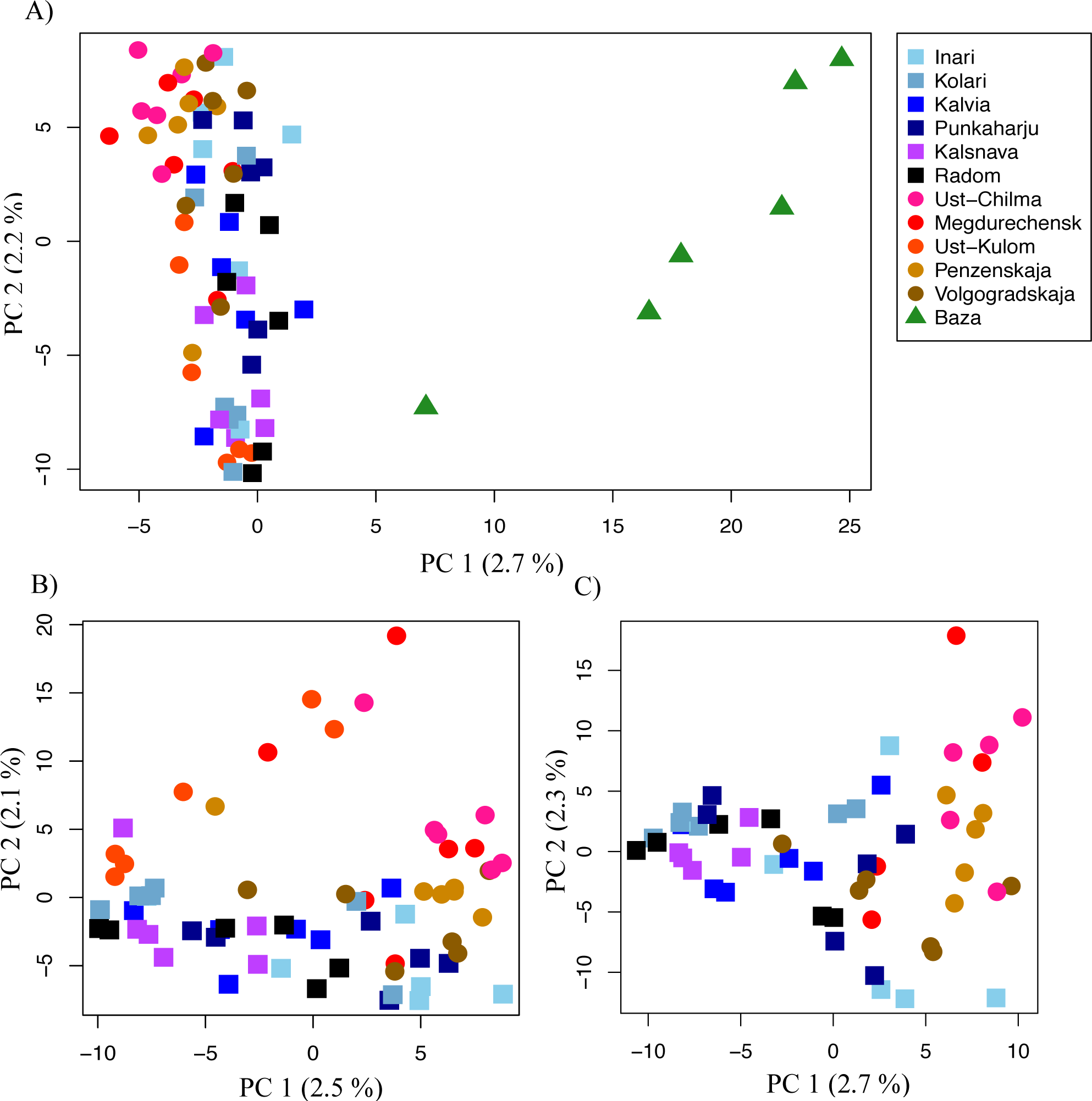
PCA projections of two first principal components of all samples (A), excluding Baza population samples (B) and excluding Baza and the samples containing the putative inversion (C). Circles represent the populations of the western cline, squares the eastern cline and triangles the isolated Baza population where included. Total variance explained by principal component is indicated within parentheses next to respective principal component axis header.

ConStruct analysis was performed using the both non-spatial model and the spatial model, which incorporated information on the geographical distance between populations. A cross-validation test indicated that for the spatial and non-spatial models the predictive accuracy improved with more layers, but only modest improvement can be seen after K2 (Figure S1). At K2 the spatial model has a better fit than the non-spatial model. Even though the K2 model has two layers, the second layer contributes very little (1-2 %) to populations other than Baza where it contributes 8 %. We also inspected the value of parameter *α_D_* which controls the shape of the decay of covariance in the spatial model, with values close to 0 indicating no isolation-by-distance (equation 3 in Bradburd et al., 2018). The first layer with larger proportion produces a value of 0.0020 for *α_D_* indicating that a very weak isolation-by-distance-pattern can be detected through most of the sampled distribution. The isolation-by-distance pattern is described by *α_D_* together with other layer parameters (*α_0_* = 0.0098, *α_2_*=0.093, *φ*=0.004, *γ*=0.144) and nugget values for each population (0.051-0.058) (Figure 5C for K1 and Figure S3 for K2). Most of the covariance shown in the figure (0.185) is explained by the covariance originating from the same ancestry (i.e. layer). The within population covariance (dots) is slightly higher by 0.06. The contribution of IBD to covariance defined by alpha-values is very small, only 0.006, but nonetheless explains the data better than a spatial model, which omits the alpha parameters.

**Figure 5.**
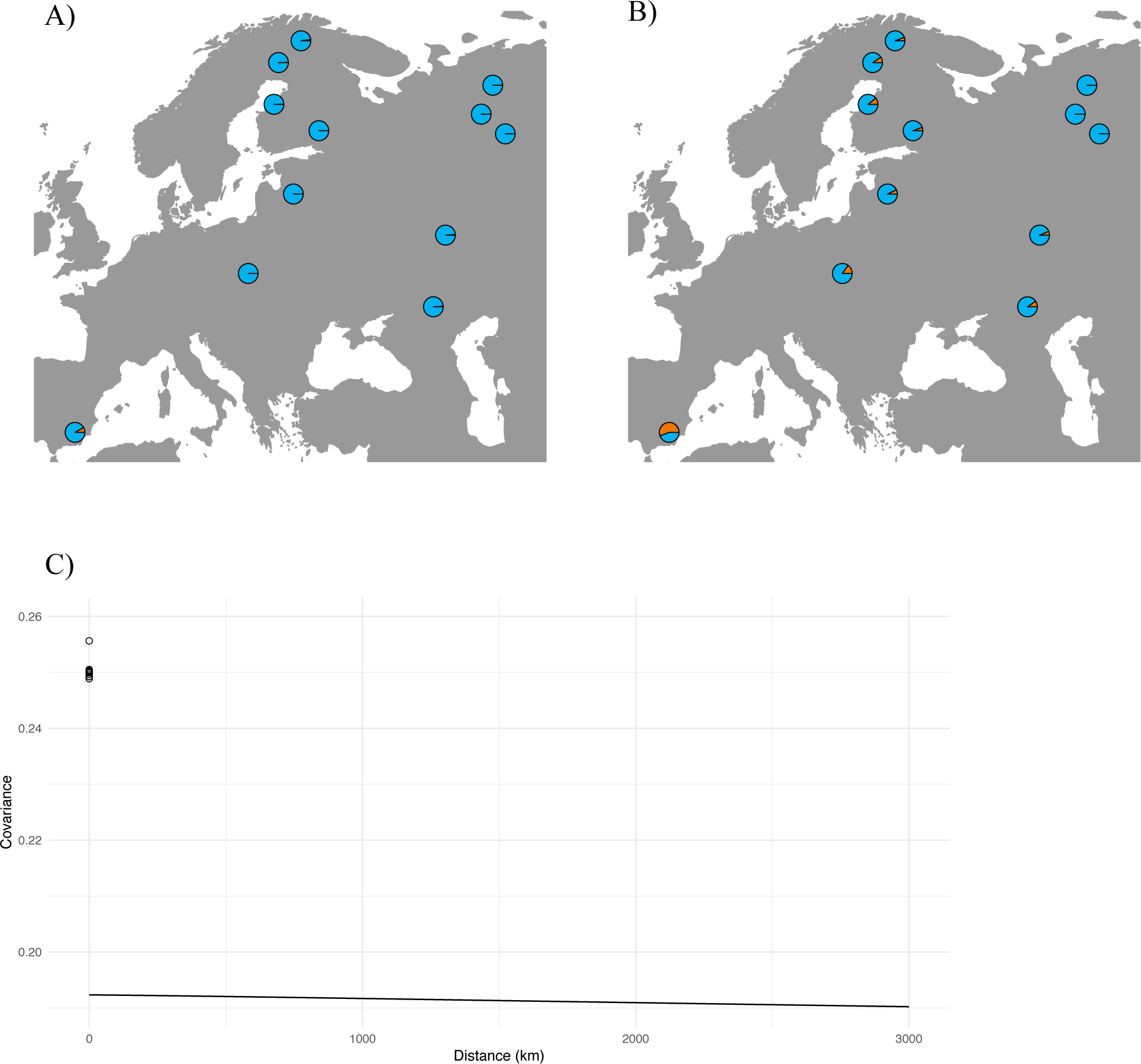
Admixture proportions for two layers estimated for different populations using conStruct A) spatial and B) non-spatial models. C) shows the within population covariance (dots) and the decrease of covariance between populations as between-population distance increases (line) for the first layer.

### Identifying loci responsible for local adaptation

#### Linear regression on latitude

Comparison of linear regression models with or without latitude was used to identify clinal allele frequency patterns. At a p-value cutoff of 0.001 a total of 263 genes were outliers, including several genes with interesting function (Table S1). However, controlling for false discovery rate (FDR, Benjamini & Hochberg, 1995) with q-values obtained from the p-value distribution no outliers at 0.10 FDR were found, suggesting that a high proportion of top candidates are false positives.

#### Genome scans for selection

Bayescan, an F_ST_ outlier detection method, was used to detect putative SNPs underlying local adaptation. Using 0.1 FDR level we obtained a single outlier locus, which is located in non-coding area of the *P. taeda* reference genome v. 1.01 in position 404,961 of tscaffold3905. However, blast search of the surrounding sequence against all known gymnosperm genes at ConGenIE (http://congenie.org/) (Sundell et al., 2015) revealed that the outlier locus appears to lie within a gene with an unknown function. The outlier locus has a distinct allele frequency pattern (Figure 6A) where the frequency of the alleles varies along latitude with steeper cline in the east. Interestingly, the second highest scoring variant, although above the 0.05 FDR limit, shows a very similar allele frequency cline pattern to the first outlier (Figure 6B). The variant is located within an intron of a Rubisco gene family member. The two top variants are not in LD nor do they appear to be located in the same scaffold of the *P. taeda* reference genome. Further, the subsequent SNPs in p-value rank, although also above the FRD limit, seem to have identical allele frequencies population-wise with high minor allele frequency especially in the Ust-Kulom population. These SNPs were further examined by studying their LD patterns.

**Figure 6.**
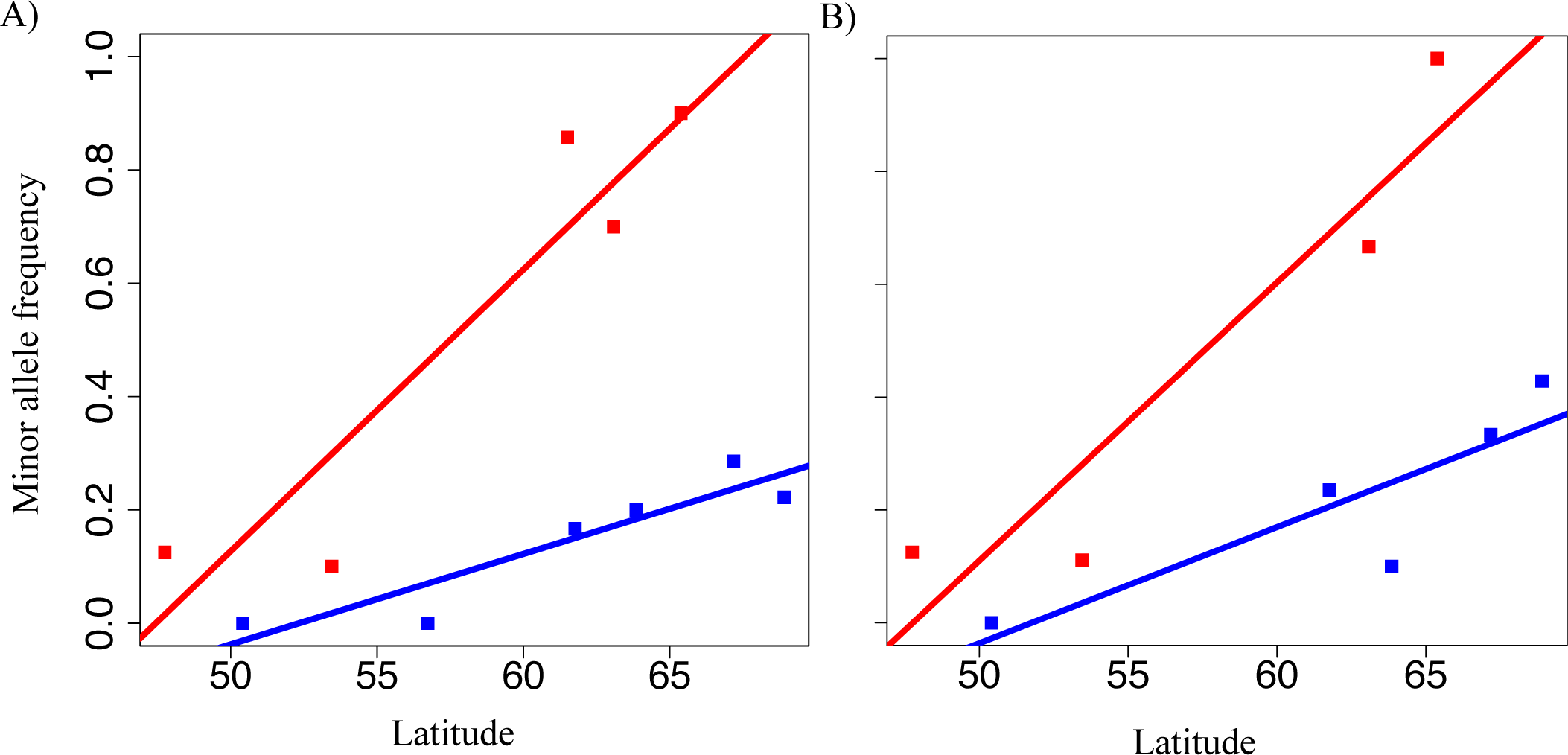
A) Bayescan outlier locus allele’s frequency at sampling sites (Y-axis) along the latitude of the sites (X-axis). Populations are marked with red (eastern) and blue (wester) squares with respective least squares trend line. B) Allele frequency of the second highest scoring bayescan result. The two figures do not share add data points due to missing data.

We also used pcadapt (Luu et al., 2017) to identify loci under selection by searching for excess divergence along principal components of population structure. P-values of each SNP assigned by pcadapt (Figure S4) were used to obtain FDR estimates. With a q-value cutoff 0.1 a total of 489 SNPs are assigned as outliers.

#### Linkage disequilibrium patterns

Linkage disequilibrium patterns (Figure 7) suggest that LD decays quickly within the *P. sylvestris* genome as the *r^2^* values fall below 0.2 within 145 bp. However, bayescan analysis revealed that many SNPs in different scaffolds had identical allele frequencies with particularly high minor allele frequency in the Ust-Kulom population. More careful examination of the LD pattern for these SNPs revealed a distinct haplotype structure in 11 samples, of which five belong to the Ust-Kulom, two to Megdurechensk, two to Penzenskaja, one to Ust-Chilma and one to Radom population.

**Figure 7.**
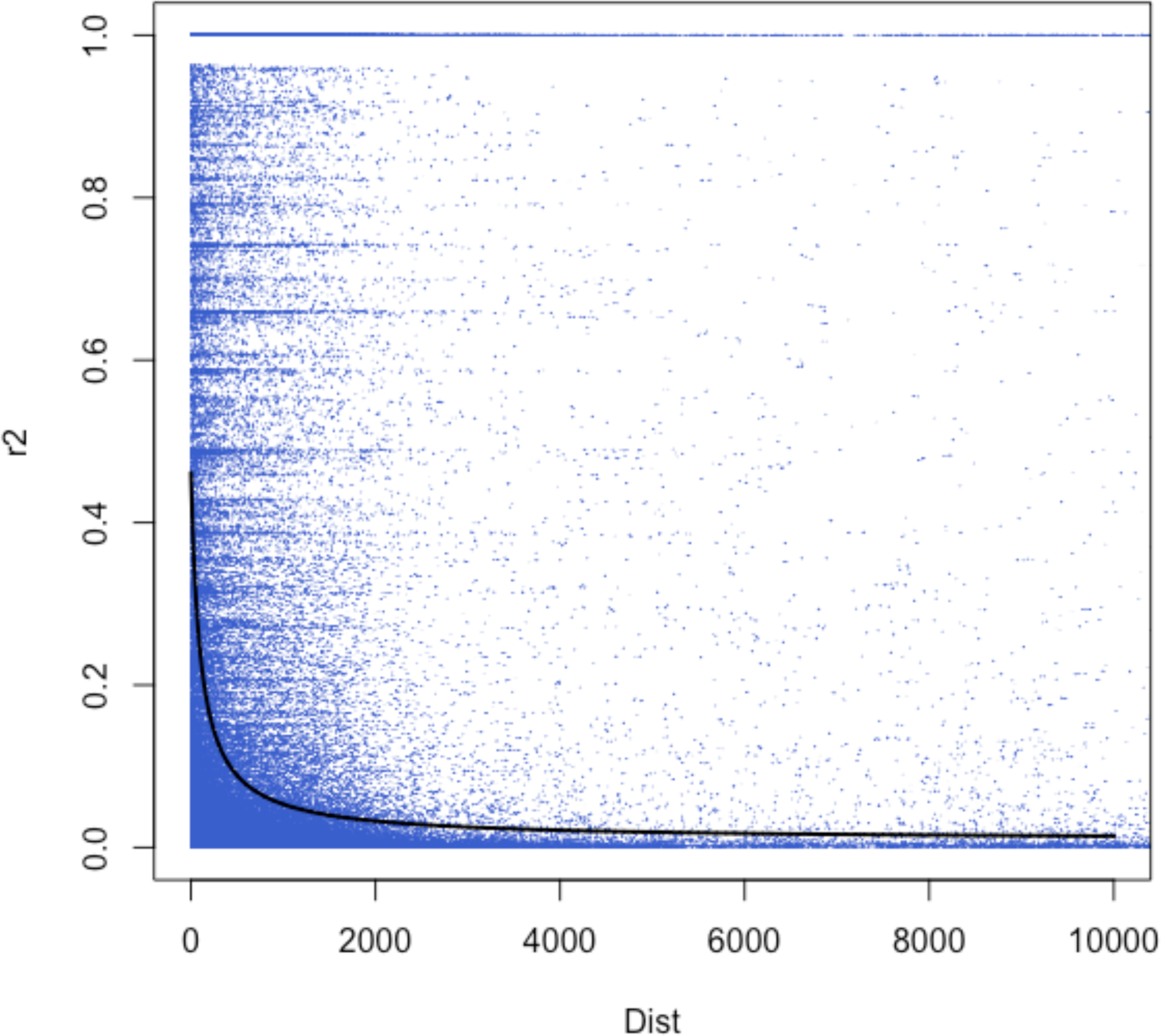
Pairwise linkage disequilibrium coefficients (r^2^) based on all pairwise SNP comparisons for all samples. Black line shows the trendline for the nonlinear regression (r^2^) against physical distance between the SNPs.

In total 169 variants located in 59 different reference sequence scaffolds had identical allele frequency and LD pattern (Figure 8A). The possibility of detecting such haplotype structure by change was explored using a permutation test showing that the chance for such is very low with a p-value of < 0.001. In the set of samples exhibiting the haplotype structure, the average nucleotide diversity within the 1-kbp area surrounding each outlier variant was only 0.0003, compared to value of 0.0034 within the same region observed between other samples. Average pairwise nucleotide diversity value calculated between the samples exhibiting the haplotype structure and other samples was 0.0093 for the same area.

**Figure 8.**
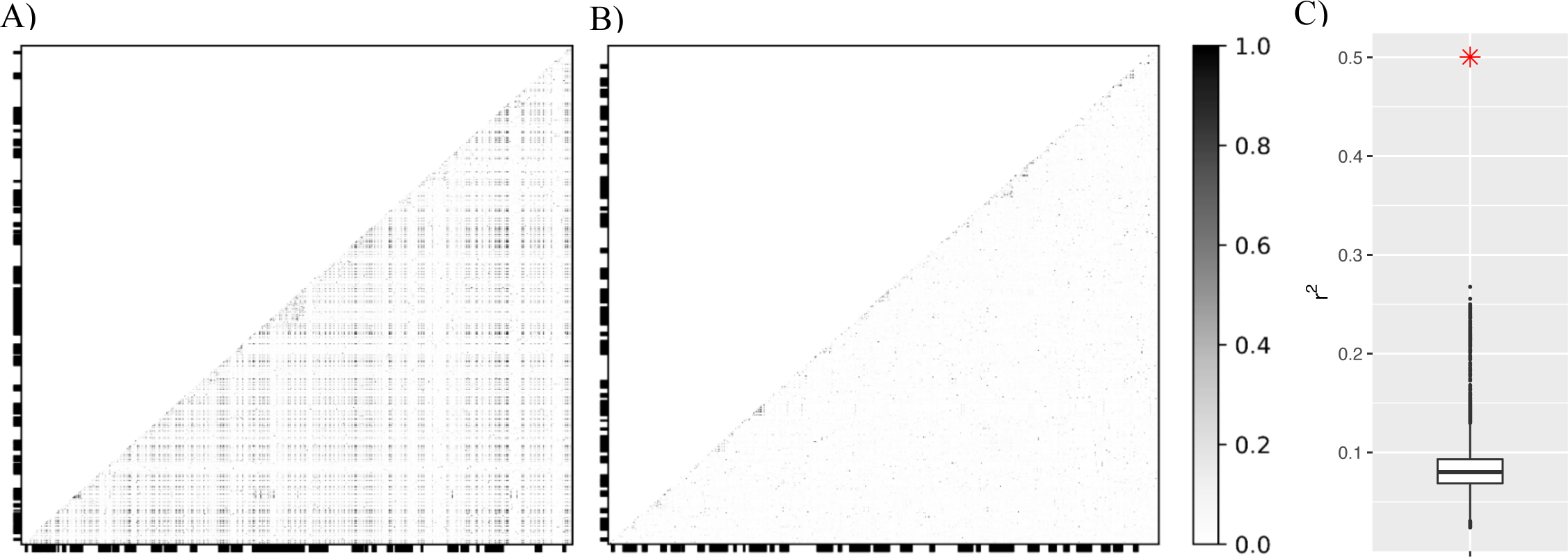
A) Heatmap visualization of allelic correlation coefficient (*r^2^*) values calculated between all SNPs identified as being part of an inversion, and all variants within their surrounding 1 kbp areas. Alternating thick and thin X and Y axis borders denotes variants belonging to the same scaffolds. B) Similar heatmap to A, but random variants with similar allele frequency to the inversion were selected along with their 1 kbp surrounding areas to visualize typical linkage disequilibrium patterns. Some scaffolds show LD within them, but between scaffolds mostly only low values *r^2^* can be seen. C) Means of *r^2^* values for 10,000 random 1kb areas (one of which is visualized in 1B heatmap) marked in black and the mean *r^2^* value of blocks containing the inversion haplotype marked in red.

Westbrook et al. (2015) generated a consensus genetic map for *P. taeda* and aligned many of their EST sequences corresponding to the marker data to the *P. taeda* reference genome v. 1.01, thus providing putative physical location for many scaffolds of the reference. There were 12 cases where the EST sequence alignment covers or is within a few kbp of the SNPs belonging to the *P. sylvestris* haplotype structure. In ten of these cases the Westbrook et al. (2015) data suggests that the scaffolds belong to linkage group one, but one scaffold seem to be part of linkage group 3 and another on of linkage group 10. The SNPs positioned in linkage group 1 are located between positions 51.12 cM and 94.54 cM. This 43.42 cM area is 23.48 % of the total length of the first linkage group. Given that the 12 chromosomes of *P. sylvestris* seem to be similar in size, we can expect each chromosome to be roughly 2.0 Gbp long. We can then, naively, take the proportion the haplotype structure covering the first chromosome’s genetic map and apply it to the expected physical size of the chromosome. This approach gives us an estimated size of 470 Mbp for the haplotype structure. Another way for estimating the haplotypes minimum size is to simply sum the lengths of the *P. taeda* reference scaffolds containing the haplotype structure, which gives a size estimate of 35.9 Mbp. As we have a capture target in 11.0 % of *P. taeda* reference genome scaffolds, we estimate that the total area covered by the haplotype structure is roughly 35.9 / 0.11 = 326 Mbp.

Allele frequencies of the haplotype structure were highly similar to the detected population structure patterns. To investigate whether the haplotype drove the observed population structure, STRUCTURE, PCA and pairwise F_ST_ analysis were redone without the scaffolds containing the haplotype structure. No change was observed in the STRUCTURE results. However, the F_ST_ results appear to be affected such that lower values are now seen between Ust-Kulom and other populations (Table S2). In the PCA results, the projection where Baza samples are omitted (Figure 5B), the samples containing the haplotype structure are separated by the second principal component and the other eastern samples are separated from western samples by the first principal component (Figure 5B). Removing also the samples that contain the haplotype structure results in projection where Eastern and Western samples occupy the opposing ends of the first principal component.

## Discussion

This first genome-wide analysis of *P. sylvestris* genetic diversity covering a large proportion of its distribution show estimates of genetic diversity and population structure largely in line with previous studies. Many loci possibly contributing to local adaptation were discovered that have been shown to be targets of selection in previous studies (Table S1) for example in *A. thaliana* (Knoth & Eulgem, 2008) and Eucalyptus (Jordan, Hoffmann, Dillon, & Prober, 2017). Furthermore, a previously unidentified large structural variation, possibly related to local adaptation, was uncovered.

### Indications of population structure

The neutral nucleotide diversity found in this study is similar to what has been observed in previous studies of this species (Kujala & Savolainen, 2012) and in several other conifers (Brown, Gill, Kuntz, Langley, & Neale, 2004; Eckert et al., 2013; Grivet et al., 2017). Given the mostly continuous distribution of wind-pollinated *P. sylvestris* and the previous findings of near-absent population structure, only very negligible differences between populations were anticipated, with the exceptions of geographically isolated populations (Kujala & Savolainen, 2012; Tanja Pyhäjärvi et al., 2007; Wachowiak, Balk, & Savolainen, 2009). It has been suggested that putative *P. sylvestris* refugia during the last glaciation has existed in the Mediterranean area, northern Europe and also in the east, possibly in Ural mountains (Naydenov et al., 2007). Therefore, genetic structure suggesting expansion from two distinct refugia, from east and west, would not have been entirely unexpected.

All approaches of population structure analysis uniformly indicated that the Baza population, geographically isolated from the main distribution, was most, although still weakly differentiated from the other populations. STRUCTURE and PCA analysis also gave some indication that the most likely number of groups is three. However, the models of these frequently used methods do not explicitly account for geographic isolation-by-distance, which can be assumed to exist within the distribution of many species, including *P. sylvestris*. Omission of this phenomenon from the models may cause these methods to spuriously assign populations to separate groups, when the genetic variation could in fact be explained by isolation-by-distance (Bradburd et al., 2018). This also seems to have happened with our *P. sylvestris* analysis, where STRUCTURE suggested three distinct clusters, but the conStruct spatial model explains the genetic covariance by within sampling location effect accompanied with weak isolation-by-distance pattern across populations.

Our results are in contrast with results from many other tree species, such as *Picea abies* where considerable structure has been detected despite many similarities in distribution, population size and reproductive biology (Chen et al., 2012). Several studies in *Populus* have also suggested the presence of distinct population structure (Evans et al., 2014; Geraldes et al., 2014; Keller et al., 2010). *P. sylvestris* rarely hybridizes with other species, and is not capable of clonal reproduction, but the exact connection between these characteristics and lack of major population structure is not understood. This lack of genome-wide structure is an advantage when investigating the genetic basis of adaptation, as such structure is a complicating factor in selection scans (Hoban et al., 2016).

### Putative signs of local adaptation

As *P. sylvestris* is known to be locally adapted to various environmental conditions within its vast distribution (Savolainen et al., 2007), we anticipated to identify signs of natural selection in the genomic variation. We performed an F_ST_-based outlier scan, which identified only single statistically significant outlier SNP with an allele frequency cline in western transect and a particularly strong cline in the eastern transect, with another non-significant variant exhibiting very similar allele frequency cline to the top outlier. The first outlier did not have any reliable annotation, but the second variant was located within an intron of a Rubisco gene, which has been suggested to have a role in ecological adaptation to different temperatures and CO_2_ concentrations (Hermida-Carrera et al., 2017). A large number of outliers in F_ST_ based selection scan would have been unexpected considering earlier findings (Kujala & Savolainen, 2012) and most theory suggesting that the nature of the underlying genetic architecture is likely highly polygenic, but observing only single outlier is surprising. It is possible that higher number of populations and samples for bayescan analysis would have allowed more outliers to be uncovered.

We also applied the *pcadapt* method, which accounts for the possible population structure via principal component analysis and identifies outliers relative to this structure. The approach yielded 489 putative outliers. As expected, the outliers also included the SNPs identified as part of the haplotype structure discussed below. Linear regression of allele frequencies to latitude was performed to discover if allele frequency clines, but even though many interesting genes were in the top candidates, no high confidence outliers were detected.

Several putative explanations exist for detecting low number of outliers in the bayescan and linear regression outlier analysis. Firstly, as the targeted sequencing approach by definition will only allow examination of very small proportion of the genome, much of the adaptive variation cannot be detected. In species with large genomes, proportionately more adaptive variation is expected to be found outside coding region (Mei, Stetter, Gates, Stitzer, & Ross-Ibarra, 2018), which may also explain the lack of adaptive signal in the data obtained by exome capture. Secondly, as discussed above, it is possible that the eastern and western parts of our sampling have in fact originated from different refugia after the most recent glacial period, or several periods as the same refugia may have existed during many or most such periods. Therefore distinctive genetic adaptations may have evolved within each refugium, as suggested for instance by Naydenov et al. (2007), rendering in particular landscape genetics approaches ineffective. Thirdly, the genetic basis of local adaptation has shown to be mostly polygenic, only a low proportion of all variants can be expected to be under strong enough selection for prolonged period to be detected. For instance Latta (1998, 2003) and Kremer and Le Corre (2012; Le Corre & Kremer, 2003b, 2012) describe how the contribution of covariance (linkage disequilibrium) between loci becomes more important relative to between population allele frequency differentiation when the number of loci contributing to genetic variation of a given trait increases. Further, Yeaman (2015) presented a model where a large number of loci with alleles of small effect contribute to local adaptation under gene flow, with temporally varying frequencies. Under such models only very limited number of large effect alleles are expected to contribute to local adaptation. As the large effect loci are often the ones detected in F_ST_-outlier tests and forming allele frequency clines, the absence of such loci in this study may not be surprising. Detection of alleles with smaller effect may require considerably larger sample size and different approaches (Berg & Coop, 2014; Field et al., 2016; Racimo, Berg, & Pickrell, 2018), but even when applying such methods it may be challenging to control for population structure to avoid false positive signal (Berg et al., 2019).

### Linkage disequilibrium patterns and putative large inversion

Earlier work has shown that LD decays very rapidly within the *P. sylvestris* genome (Dvornyk, Sirviö, Mikkonen, & Savolainen, 2002; Tanja Pyhäjärvi et al., 2007; Wachowiak et al., 2009) and also in other conifers such as *P. taeda* (Acosta et al., 2019; Lu et al., 2016). This study allows for examining longer range patterns of LD than before as in many cases multiple target sequences are positioned within the same scaffold. Our findings are in line with the previous studies showing that *r^2^* fall below 0.2 within 145 bp. The advantage of low the level of genome-wide LD observed previously (Tanja Pyhäjärvi et al., 2007), and in this work, is that variants detected in the outlier scans are probably very close to the causative polymorphism (Neale & Savolainen, 2004).

Interestingly, the bayescan analysis revealed a large number of SNPs forming an unexpected haplotype structure. Given that the LD decays very quickly within the *P. sylvestris* genome, the LD pattern detected here is very likely not due to chance. Analysis of pairwise nucleotide differences of the affected region shows that very low level of nucleotide diversity can be observed within the samples where the haplotype is present, but in the other samples the diversity level seems comparable to average genome-wide level. Due to the fragmented nature of the *P. taeda* reference genome, and the lack of *P. sylvestris* reference, it is not possible to directly verify the position and the size of the haplotype structure, but some crude estimates could be made to indicate that it is very likely to be several hundred million base pairs long.

We did not find any putative technical explanations for the phenomenon, as stringent parameters were used in the read alignment and SNP calling, where only concordantly, uniquely aligned reads were retained. In addition, the read alignments were visually inspected using samtools tview, and no abnormalities were found. Haplotype patterns can be created by partial selective sweeps but considering that the haplotype structure spans 43 cM in the *P. taeda* genetic map, this explanation of sweep does not seem possible under the normal recombination rates, but requires structural variation. Therefore, inversion considerably reducing recombination and extending impact of partial sweep is a very likely explanation for the phenomenon.

Possible role of inversions in local adaptation had been recognized in genetics research early on (Dobzhansky, 1970), and the concept of ‘supergenes’ has since been further explored (Thompson & Jiggins, 2014). Kirkpatrick and Barton (2006) presented models to study how inversions containing beneficial variants may contribute to local adaptation in various scenarios. They show that if an inversion encompasses two or more locally adapted alleles, it has a fitness advantage over other haplotypes with fewer locally adapted alleles. A further contribution to the fitness advantage may occur, if the inversion contains alleles with positive epistasis (Feldman, Otto, & Christiansen, 1997). Also, particularly low deleterious mutation load within the inversion can lead or contribute to fitness advantage (Nei, Kojima, & Schaffer, 1967). If fitness advantage arises, the inversion will rise in frequency within the geographic area it provides selective advantage until it reaches migration-selection balance. Locally advantageous mutations can also accumulate after initial spread (Faria, Johannesson, Butlin, & Westram, 2019). Inversions can create large areas of restricted recombination as they prevent proper chromatid pairing, although gene conversion and double cross-overs may allow some genetic exchange between inverted and non-inverted haplotypes (Andoflatto, Depaulis, & Navarro, 2001). Empirical studies have inversions contributing to local adaptation for instance in *D. melanogaster* (Kapun & Flatt, 2018), sticklebacks (Jones et al., 2012), yellow monkeyflower (Gould, Chen, & Lowry, 2018), teosinte (Tanja Pyhäjärvi, Hufford, Mezmouk, & Ross-Ibarra, 2013), humans (Puig, Casillas, Villatoro, & Cáceres, 2015) and in many others (Wellenreuther & Bernatchez, 2018).

Pine genomes typically have high synteny even between species (Komulainen et al., 2003). Signs of putative inversions within *P. sylvestris* are rare in literature, although some earlier cytological studies have detected putative inversions in the eastern part of the species distribution (Muratova, 1994), but many observations of inversions have been made in other conifer species (e.g. Pederick, 1968). The reason relatively few large inversions have been detected may be due to difficulty in detecting them, as contemporary resequencing or genome assembly approaches combined even with the current long-read sequencing platforms may not be sufficient, and often complementary approaches such as optical mapping are required (Kronenberg et al., 2018; Levy-Sakin et al., 2019). The detection may be particularly difficult in conifers due to the fragmented nature of their reference genomes, as common methods for detecting of structural variation often rely on examining long scaffolds or identifying exact breakpoints making targeted sequencing data unsuitable (Cáceres & González, 2015).

The size of the putative inversion found here is very large, although sizeable inversions, up to 191 Mbp, have currently been predicted in the human genome in the invFEST database (Martínez-Fundichely et al., 2014). Large inversions have been suggested as being targets of selection in many species (Wellenreuther & Bernatchez, 2018), with the largest such inversions exceeding 200 Mbp between two *Helianthus* sister species (Barb et al., 2014). To our knowledge, the putative inversion detected in our study remains as the largest one to suggested to contribute to local adaptation to date, although the relative size the inversion compared to whole genome size is certainly not as large as inversions in some other species. If more such large inversions exist in the *P. sylvestris* genome, it can also have drastic effects for breeding efforts as recombination is very limited within the inversion in polymorphic crosses.

If the inversion contains loci with alternative alleles affecting the timing of bud set formation and similar traits contributing to adaptation to environmental gradients, selection for local adaptation may have driven the inversion to its current high local frequencies. The samples containing the inversion polymorphism seem to be located broadly in the similar, but not identical, geographic area where STRUCTURE and PCA analysis indicate slight population structure border. This may suggest that the inversion event may have occurred within the speculated eastern refugium, but further investigation would be required to uncover the possible origin and fitness effects. Reflecting the suggestion of large inversions contributing to reproductive isolation (Noor, Gratos, Bertucci, & Reiland, 2001) to our findings of samples containing the putative inversion appearing somewhat differentiated in PCA and the respective populations in F_ST_ results, raises the question whether barrier to gene exchange is now introduced between samples containing the inversion and others. It is difficult to rule out such hypothesis without crossing experiments, but there is no evidence clearly suggesting this. In the case of PCA the lack of any other structure within the main species range may lead to projections emphasizing the difference within the inversion. F_ST_ results between Ust-Kulom, where the inversion has highest frequency, and others are reduced when scaffolds known to be part of the inversion are removed, although they still remain elevated. It may be that several scaffolds that are part of the inversion are not identified as part of it, as they happen to not contain the SNPs used to identify the inversion. STRUCTURE and conStruct results remain unaffected by the removal of the putative inversion.

## Conclusions

In this work we have examined the genetic diversity of *P. sylvestris* along a large portion of its range. Some patterns of population structure can be seen in a marginal population but within the continuous main range the isolation-by-distance explains well any differentiation detected, unlike in many other tree species. This mitigates the issues caused by structure in detecting signs of selection, but our results also show that while clear phenotypic signals of local adaptation have been detected, the molecular background remains largely elusive even if many well-established approaches were used here to detect the signature of selection. However, many interesting outliers were detected that have been shown to contribute to local adaptation in earlier studies. Furthermore, in this study we find a putative very large inversion, likely spanning an area equivalent to several *Arabidopsis thaliana* genomes. To our knowledge, this is the first time that an inversion has been shown to putatively contribute to local adaptation in conifers, even though such occurrences can certainly be expected by theory (Yeaman, 2013).

## Acknowledgements

The authors thank Skogforsk for providing seeds for sequencing, Matias Kirst for helping with developing the exome capture protocol, Gideon Bradburd for help with conStruct and members of the Plant Genetics Research group in the University of Oulu for many helpful comments and suggestions. We thank the CSC-IT Center for Science, Finland, for computational resources. This work was supported by European Comission 7^th^ Framework Programme project ProCoGen (289841) to O.S., Biocenter Oulu, Emil Aaltosen Säätiö (160284 O), Oulun Läänin Talousseuran Maataloussäätiö to JT, Academy of Finland (287431 and 293819) to T.P. Z.L. is funded by a postdoctoral fellowship from the Special Research Fund of Ghent University (BOFPDO2018001701).

## Author Contributions

J.S.T., O.S. and T.P. designed the study. J.S.T., Z.L., L.S., J.J.A, J.V. and M.T.C. generated data and analytical tools. J.S.T. and T.P. analyzed the data and wrote the manuscript with O.S. All authors commented the manuscript.

## References

Acosta, J. J., Fahrenkrog, A. M., Neves, L. G., Resende, M. F. R., Dervinis, C., Davis, J. M., … Kirst, M. (2019). Exome Resequencing Reveals Evolutionary History, Genomic Diversity, and Targets of Selection in the Conifers Pinus taeda and Pinus elliottii. Genome Biology and Evolution, 11(2), 508–520.

Adrion, J. R., Hahn, M. W., & Cooper, B. S. (2015). Revisiting classic clines in Drosophila melanogaster in the age of genomics. Trends in Genetics, 31(8), 434–444. doi:10.1016/j.tig.2015.05.006

Ågren, J., & Schemske, D. W. (2012). Reciprocal transplants demonstrate strong adaptive differentiation of the model organism Arabidopsis thaliana in its native range. New Phytologist, 194(4), 1112–1122.

Aho, M. (1994). Autumn frost hardening of one-year-old *Pinus sylvestris* (L.) seedlings: Effect of origin and parent trees. Scandinavian Journal of Forest Research, 9(1–4), 17–24. doi:10.1080/02827589409382808

Alberto, F. J., Aitken, S. N., Alía, R., González-Martínez, S. C., Hänninen, H., Kremer, A., … Savolainen, O. (2013). Potential for evolutionary responses to climate change - evidence from tree populations. Global Change Biology, 19(6), 1645–61. doi:10.1111/gcb.12181

Alberto, F. J., Derory, J., Boury, C., Frigerio, J.-M., Zimmermann, N. E., & Kremer, A. (2013). Imprints of natural selection along environmental gradients in phenology-related genes of Quercus petraea. Genetics, 195(2), 495–512. doi:10.1534/genetics.113.153783

Alexander, D. H., Novembre, J., & Lange, K. (2009). Fast model-based estimation of ancestry in unrelated individuals, 1655–1664. doi:10.1101/gr.094052.109.vidual

Andoflatto, P., Depaulis, F., & Navarro, A. (2001). Inversion polymorphisms and nucleotide variability in Drosophila. Genetics Research, 77(01), 1–8. doi:10.1017/S0016672301004955

Barb, J. G., Bowers, J. E., Renaut, S., Rey, J. I., Knapp, S. J., Rieseberg, L. H., & Burke, J. M. (2014). Chromosomal evolution and patterns of introgression in Helianthus. Genetics, 197(3), 969–979. doi:10.1534/genetics.114.165548

Barton, N. H. (1999). Clines in polygenic traits. Genetical Research, 74(3), S001667239900422X. doi:10.1017/S001667239900422X

Barton, N. H., & Keightley, P. D. (2002). Multifactorial Geneticsunderstanding Quantitative Genetic Variation. Nature Reviews Genetics, 3(1), 11–21. doi:10.1038/nrg700

Benjamini, Y., & Hochberg, Y. (1995). Controlling the False Discovery Rate: A Practical and Powerful Approach to Multiple Testing. Journal of the Royal Statistical Society. Series B (Methodological). WileyRoyal Statistical Society. doi:10.2307/2346101

Berg, J. J., & Coop, G. (2014). A Population Genetic Signal of Polygenic Adaptation. PLoS Genetics, 10(8). doi:10.1371/journal.pgen.1004412

Berg, J. J., Harpak, A., Sinnott-Armstrong, N., Joergensen, A. M., Mostafavi, H., Field, Y., … others. (2019). Reduced signal for polygenic adaptation of height in UK Biobank. ELife, 8, e39725.

Beuker, E. (1994). Adaptation to climatic changes of the timing of bud burst in populations of Pinus sylvestris L. and Picea abies. Tree Physiology, 14(7–9), 961–970.

Bhatia, G., Patterson, N., Sankararaman, S., & Price, A. L. (2013). Estimating and interpreting FST: The impact of rare variants. Genome Research, 23(9), 1514–1521. doi:10.1101/gr.154831.113

Bradburd, G. S., Coop, G. M., & Ralph, P. L. (2018). Inferring continuous and discrete population genetic structure across space. Genetics, 210(1), 33–52.

Brown, G. R., Gill, G. P., Kuntz, R. J., Langley, C. H., & Neale, D. B. (2004). Nucleotide diversity and linkage disequilibrium in loblolly pine. Proceedings of the National Academy of Sciences of the United States of America, 101(42), 15255–15260. doi:10.1073/pnas.0404231101

Buckler, E. S., Holland, J. B., Bradbury, P. J., Acharya, C. B., Brown, P. J., Browne, C., … others. (2009). The genetic architecture of maize flowering time. Science, 325(5941), 714–718.

Cáceres, A., & González, J. R. (2015). Following the footprints of polymorphic inversions on SNP data: From detection to association tests. Nucleic Acids Research, 43(8). doi:10.1093/nar/gkv073

Cheddadi, R., Vendramin, G. G., Litt, T., François, L., Kageyama, M., Lorentz, S., … Lunt, D. (2006). Imprints of glacial refugia in the modern genetic diversity of Pinus sylvestris. Global Ecology and Biogeography, 15(3), 271–282. doi:10.1111/j.1466-822X.2006.00226.x

Chen, J., Källman, T., Ma, X., Gyllenstrand, N., Zaina, G., Morgante, M., … Lascoux, M. (2012). Disentangling the roles of history and local selection in shaping clinal variation of allele frequencies and gene expression in Norway spruce (Picea abies). Genetics, 191(3), 865–81. doi:10.1534/genetics.112.140749

Coop, G., Witonsky, D., Di Rienzo, A., & Pritchard, J. K. (2010). Using environmental correlations to identify loci underlying local adaptation. Genetics, 185(4), 1411–23. doi:10.1534/genetics.110.114819

Danecek, P., Auton, A., Abecasis, G., Albers, C. A., Banks, E., DePristo, M. A., … Durbin, R. (2011). The variant call format and VCFtools. Bioinformatics (Oxford, England), 27(15), 2156–8. doi:10.1093/bioinformatics/btr330

Dobzhansky, T. (1970). Genetics of the evolutionary process (Vol. 139). Columbia University Press.

Dvornyk, V., Sirviö, A., Mikkonen, M., & Savolainen, O. (2002). Low nucleotide diversity at the pal1 locus in the widely distributed Pinus sylvestris. Molecular Biology and Evolution, 19(2), 179–188. doi:10.1093/oxfordjournals.molbev.a004070

Eckert, A. J., Bower, A. D., Jermstad, K. D., Wegrzyn, J. L., Knaus, B. J., Syring, J. V., & Neale, D. B. (2013). Multilocus analyses reveal little evidence for lineage-wide adaptive evolution within major clades of soft pines (Pinus subgenus Strobus). Molecular Ecology, 22(22), 5635–5650. doi:10.1111/mec.12514

Eiche, V. (1966). Cold Damage and Plant Mortality in Experimental Provenance Plantations with Scots Pine in Northern Sweden. STUDIA FORESTALIA SUECICA, 36, 1–219.

Evanno, G., Regnaut, S., & Goudet, J. (2005). Detecting the number of clusters of individuals using the software STRUCTURE: A simulation study. Molecular Ecology, 14(8), 2611– 2620. doi:10.1111/j.1365-294X.2005.02553.x

Evans, L. M., Slavov, G. T., Rodgers-Melnick, E., Martin, J., Ranjan, P., Muchero, W., … DiFazio, S. P. (2014). Population genomics of Populus trichocarpa identifies signatures of selection and adaptive trait associations. Nature Genetics, 46(10), 1089–1096. doi:10.1038/ng.3075

Excoffier, L., Hofer, T., & Foll, M. (2009). Detecting loci under selection in a hierarchically structured population. Heredity, 103(4), 285–98. doi:10.1038/hdy.2009.74

Fan, S., Hansen, M. E. B., Lo, Y., & Tishkoff, S. A. (2016). Review of Recent Human Adaptation. Science, 354(6308), 54–59. doi:10.1126/science.aaf5098

Faria, R., Johannesson, K., Butlin, R. K., & Westram, A. M. (2019). Evolving Inversions. Trends in Ecology and Evolution, 34(3), 239–248. doi:10.1016/j.tree.2018.12.005

Feldman, M. W., Otto, S. P., & Christiansen, F. B. (1997). Population Genetic Perspectives on the Evolution of Recombination. Annual Review of Genetics, 30(1), 261–295. doi:10.1146/annurev.genet.30.1.261

Field, Y., Boyle, E. A., Telis, N., Gao, Z., Gaulton, K. J., Golan, D., … Pritchard, J. K. (2016). Detection of human adaptation during the past 2000 years. Science, 354(6313), 760–764. doi:10.1126/science.aag0776

Fisher, R. (1918). The Correlation Between Relatives on the Supposition of Mendelian Inheritance. Proc. Roy. Soc. Edinburgh, 52, 399–433.

Foll, M., & Gaggiotti, O. (2008). A genome-scan method to identify selected loci appropriate for both dominant and codominant markers: a Bayesian perspective. Genetics, 180(2), 977–93. doi:10.1534/genetics.108.092221

Galinsky, K. J., Bhatia, G., Loh, P. R., Georgiev, S., Mukherjee, S., Patterson, N. J., & Price, A. L. (2016). Fast Principal-Component Analysis Reveals Convergent Evolution of ADH1B in Europe and East Asia. American Journal of Human Genetics, 98(3), 456–472. doi:10.1016/j.ajhg.2015.12.022

Garner, A. G., Kenney, A. M., Fishman, L., & Sweigart, A. L. (2016). Genetic loci with parent-of-origin effects cause hybrid seed lethality in crosses between Mimulus species. New Phytologist, 211(1), 319–331. doi:10.1111/nph.13897

Geraldes, A., Farzaneh, N., Grassa, C. J., McKown, A. D., Guy, R. D., Mansfield, S. D., … Cronk, Q. C. B. (2014). Landscape genomics of Populus trichocarpa: the role of hybridization, limited gene flow, and natural selection in shaping patterns of population structure. Evolution, 68(11), 3260–3280.

Gould, B. A., Chen, Y., & Lowry, D. B. (2018). Gene Regulatory Divergence Between Locally Adapted Ecotypes in Their Native Habitats. Molecular Ecology, (January), 1–15. doi:10.1111/mec.14852

Grivet, D., Avia, K., Vaattovaara, A., Eckert, A. J., Neale, D. B., Savolainen, O., & González-Martínez, S. C. (2017). High rate of adaptive evolution in two widespread European pines. Molecular Ecology, 26(24), 6857–6870. doi:10.1111/mec.14402

Gutenkunst, R. N., Hernandez, R. D., Williamson, S. H., & Bustamante, C. D. (2009). Inferring the joint demographic history of multiple populations from multidimensional SNP frequency data. PLoS Genetics, 5(10). doi:10.1371/journal.pgen.1000695

Hämälä, T., Mattila, T. M., & Savolainen, O. (2018). Local adaptation and ecological differentiation under selection, migration, and drift in Arabidopsis lyrata*. Evolution, 72(7), 1373–1386. doi:10.1111/evo.13502

Hermida-Carrera, C., Fares, M. A., Fernández, Á., Gil-Pelegrín, E., Kapralov, M. V, Mir, A., … others. (2017). Positively selected amino acid replacements within the RuBisCO enzyme of oak trees are associated with ecological adaptations. PloS One, 12(8), e0183970.

Hill, W. G., & Robertson, A. (1968). Linkage disequilibrium in finite populations. TAG Theoretical and Applied Genetics, 38(6), 226–231. doi:10.1007/bf01245622

Hoban, S., Kelley, J. L., Lotterhos, K. E., Antolin, M. F., Bradburd, G., Lowry, D. B., … Whitlock, M. C. (2016). Finding the Genomic Basis of Local Adaptation: Pitfalls, Practical Solutions, and Future Directions. The American Naturalist, 188(4), 379–397. doi:10.1086/688018

Holliday, J. a., Ritland, K., & Aitken, S. N. (2010). Widespread, ecologically relevant genetic markers developed from association mapping of climate-related traits in Sitka spruce (Picea sitchensis). New Phytologist, 188(2), 501–514.

Hudson, R. R., Slatkin, M., & Maddison, W. P. (1992). Estimation of levels of gene flow from DNA sequence data. Genetics, 132(2).

Hurme, P., Repo, T., Savolainen, O., & Pääkkönen, T. (1997). Climatic adaptation of bud set and frost hardiness in Scots pine (*Pinus sylvestris*). Canadian Journal of Forest Research, 27(5), 716–723. doi:10.1139/x97-052

Hurme, P., Sillanpää, M., Repo, T., Arjas, E., & Savolainen, O. (2000). Genetic basis of climatic adaptation in Scots pine by Bayesian QTL analysis. Genetics, 156(1930), 1309–1322.

Huxley, J. (1938). Clines: an Auxiliary Taxonomic Principle. Nature, 142, 219–220. doi:10.1038/142219a0

Jones, F. C., Grabherr, M. G., Chan, Y. F., Russell, P., Mauceli, E., Johnson, J., … Kingsley, D. M. (2012). The genomic basis of adaptive evolution in threespine sticklebacks. Nature, 484(7392), 55–61. doi:10.1038/nature10944

Jordan, R., Hoffmann, A. A., Dillon, S. K., & Prober, S. M. (2017). Evidence of genomic adaptation to climate in Eucalyptus microcarpa: implications for adaptive potential to projected climate change. Molecular Ecology, 0–2. doi:10.1111/mec.14341

Kapun, M., & Flatt, T. (2018). The adaptive significance of chromosomal inversion polymorphisms in Drosophila melanogaster. Molecular Ecology. doi:10.1111/mec.14871

Karhu, A., Hurme, P., Karjalainen, M., Karvonen, P., Kärkkäinen, K., Neale, D., & Savolainen, O. (1996). Do molecular markers reflect patterns of differentiation in adaptive traits of conifers? Theoretical and Applied Genetics, 93(1–2), 215–221. doi:10.1007/s001220050268

Kawecki, T. J., & Ebert, D. (2004). Conceptual issues in local adaptation. Ecology Letters, 7(12), 1225–1241. doi:10.1111/j.1461-0248.2004.00684.x

Keller, S. R., Olson, M. S., Salim, S., William, S. A., & Peter, T. (2010). Genomic diversity, population structure, and migration following rapid range expansion in the Balsam Poplar, Populus balsamifera. Molecular Ecology, 19(6), 1212–1226. doi:10.1111/j.1365-294X.2010.04546.x

Kirkpatrick, M., & Barton, N. (2006). Chromosome Inversions, Local Adaptation and Speciation, 434(May), 419–434. doi:10.1534/genetics.105.047985

Knoth, C., & Eulgem, T. (2008). The oomycete response gene LURP1 is required for defense against Hyaloperonospora parasitica in Arabidopsis thaliana. The Plant Journal, 55(1), 53–64.

Komulainen, P., Brown, G. R., Mikkonen, M., Karhu, A., Garc??a-Gil, M. R., O’Malley, D., … Savolainen, O. (2003). Comparing EST-based genetic maps between Pinus sylvestris and Pinus taeda. Theoretical and Applied Genetics, 107(4), 667–678. doi:10.1007/s00122-003-1312-2

Kopelman, N. M., Mayzel, J., Jakobsson, M., Rosenberg, N. A., & Mayrose, I. (2015). Clumpak: A program for identifying clustering modes and packaging population structure inferences across K. Molecular Ecology Resources, 15(5), 1179–1191. doi:10.1111/1755-0998.12387

Kremer, A., & Le Corre, V. (2012). Decoupling of differentiation between traits and their underlying genes in response to divergent selection. Heredity, 108(4), 375–385. doi:10.1038/hdy.2011.81

Kronenberg, Z. N., Fiddes, I. T., Gordon, D., Murali, S., Cantsilieris, S., Meyerson, O. S., … Eichler, E. E. (2018). High-resolution comparative analysis of great ape genomes. Science, 360(6393). doi:10.1126/science.aar6343

Kujala, S., Knürr, T., Kärkkäinen, K., Neale, D. B., Sillanpää, M. J., & Savolainen, O. (2017). Genetic heterogeneity underlying variation in a locally adaptive clinal trait in Pinus sylvestris revealed by a Bayesian multipopulation analysis. Heredity, 118(5), 413–423. doi:10.1038/hdy.2016.115

Kujala, S., & Savolainen, O. (2012). Sequence variation patterns along a latitudinal cline in Scots pine (Pinus sylvestris): Signs of clinal adaptation? Tree Genetics and Genomes, 8(6), 1451–1467. doi:10.1007/s11295-012-0532-5

Langmead, B., & Salzberg, S. L. (2012). Fast gapped-read alignment with Bowtie 2. Nature Methods, 9(4), 357–9. doi:10.1038/nmeth.1923

Latta, R. G. (1998). Differentiation of Allelic Frequencies at Quantitative Trait Loci Affecting Locally Adaptive Traits Differentiation of Allelic Frequencies at Quantitative Trait Loci Affecting Locally Adaptive Traits. The American Naturalist, 151(3), 283–292. doi:10.1086/286119

Latta, R. G. (2003). Gene flow, adaptive population divergence and comparative population structure across loci. New Phytologist, 161(1), 51–58. doi:10.1046/j.1469-8137.2003.00920.x

Le Corre, V., & Kremer, A. (2003a). Genetic variability at neutral markers, quantitative trait loci and trait in a subdivided population under selection. Genetics, 164(3), 1205–1219. Retrieved from http://www.scopus.com/inward/record.url?eid=2-s2.0-0043210649&partnerID=tZOtx3y1

Le Corre, V., & Kremer, A. (2003b). Genetic variability at neutral markers, quantitative trait loci and trait in a subdivided population under selection. Genetics, 164(3), 1205–1219. doi:10.1186/1471-2148-9-177

Le Corre, V., & Kremer, A. (2012). The genetic differentiation at quantitative trait loci under local adaptation. Molecular Ecology, 21(7), 1548–66. doi:10.1111/j.1365-294X.2012.05479.x

Leinonen, P. H., Remington, D. L., & Savolainen, O. (2011). Local adaptation, phenotypic differentiation, and hybrid fitness in diverged natural populations of Arabidopsis lyrata. Evolution; International Journal of Organic Evolution, 65(1), 90–107. doi:10.1111/j.1558-5646.2010.01119.x

Levy-Sakin, M., Pastor, S., Mostovoy, Y., Li, L., Leung, A. K. Y., McCaffrey, J., … Kwok, P. Y. (2019). Genome maps across 26 human populations reveal population-specific patterns of structural variation. Nature Communications, 10(1), 1–14. doi:10.1038/s41467-019-08992-7

Lewontin, R. C., & Krakauer, J. (1973). Distribution of gene frequency as a test of the theory of the selective neutrality of polymorphisms. Genetics, 74(1), 175–195.

Li, H., Handsaker, B., Wysoker, A., Fennell, T., Ruan, J., Homer, N., … Durbin, R. (2009). The Sequence Alignment/Map format and SAMtools. Bioinformatics, 25(16), 2078–2079. doi:10.1093/bioinformatics/btp352

Li, Z., De La Torre, A. R., Sterck, L., Cánovas, F. M., Avila, C., Merino, I., … Van De Peer, Y. (2017). Single-copy genes as molecularmarkers for phylogenomic studies in seed plants. Genome Biology and Evolution, 9(5), 1130–1147. doi:10.1093/gbe/evx070

Lu, M., Krutovsky, K. V, Nelson, C. D., Koralewski, T. E., Byram, T. D., & Loopstra, C. A. (2016). Exome genotyping, linkage disequilibrium and population structure in loblolly pine (Pinus taeda L.). BMC Genomics, 1–11. doi:10.1186/s12864-016-3081-8

Luu, K., Bazin, E., & Blum, M. G. B. (2017). pcadapt: an R package to perform genome scans for selection based on principal component analysis. Molecular Ecology Resources, 17(1), 67–77. doi:10.1111/1755-0998.12592

Ma, X. F., Hall, D., St. Onge, K. R., Jansson, S., & Ingvarsson, P. K. (2010). Genetic differentiation, clinal variation and phenotypic associations with growth cessation across the Populus tremula photoperiodic pathway. Genetics, 186(3), 1033–1044. doi:10.1534/genetics.110.120873

Martínez-Fundichely, A., Casillas, S., Egea, R., Ràmia, M., Barbadilla, A., Pantano, L., … Cáceres, M. (2014). InvFEST, a database integrating information of polymorphic inversions in the human genome. Nucleic Acids Research, 42(D1), 1027–1032. doi:10.1093/nar/gkt1122

McVean, G. (2009). A genealogical interpretation of principal components analysis. PLoS Genetics, 5(10). doi:10.1371/journal.pgen.1000686

Mei, W., Stetter, M. G., Gates, D. J., Stitzer, M. C., & Ross-Ibarra, J. (2018). Adaptation in plant genomes: Bigger is different. American Journal of Botany, 105(1), 16–19. doi:10.1002/ajb2.1002

Mikola, J. (1982). Bud-set phenology as an indicator of climatic adaptation of Scots pine in Finland [Pinus sylvestris]. Population Genetics of Forest Trees, 16(2), 178–184.

Muratova, E. N. (1994). Cytogenetical study on Scots pine (Pinus sylvestris L.) in the Central Yakutia. Cytogenetic Studies of Forest Trees and Shrub Species: Proc. First IUFRO Cytogenetics Working Party, 2–04. Retrieved from http://18a.akadem.ru/Articles/LSVD/2/Muratova4.pdf

Naydenov, K., Senneville, S., Beaulieu, J., Tremblay, F., & Bousquet, J. (2007). Glacial vicariance in Eurasia: Mitochondrial DNA evidence from Scots pine for a complex heritage involving genetically distinct refugia at mid-northern latitudes and in Asia Minor. BMC Evolutionary Biology, 7(1), 1–12. doi:10.1186/1471-2148-7-233

Neale, D. B., & Savolainen, O. (2004). Association genetics of complex traits in conifers. Trends in Plant Science, 9(7), 325–330.

Neale, D. B., Wegrzyn, J. L., Stevens, K. A., Zimin, A. V, Puiu, D., Crepeau, M. W., … Langley, C. H. (2014). Decoding the massive genome of loblolly pine using haploid DNA and novel assembly strategies. Genome Biology, 15(3), R59. doi:10.1186/gb-2014-15-3-r59

Nei, M., Kojima, K.-I., & Schaffer, H. E. (1967). Frequency changes of new inversions in populations under mutation-selection equilibria. Genetics, 57(4), 741.

Nei, M., & Li, W.-H. (1979). Mathematical model for studying genetic variation in terms of restriction endonucleases, 76(10), 5269–5273.

Noor, M. A. F., Gratos, K. L., Bertucci, L. A., & Reiland, J. (2001). Chromosomal inversions and the reproductive isolation of species. Proceedings of the National Academy of Sciences of the United States of America, 98(21), 12084–12088. doi:10.1073/pnas.221274498

Orr, H. A. (1998). The population genetics of adaptation: The distribution of factors fixed during adaptive evolution. Evolution, 52(4), 935–949. Retrieved from http://www.scopus.com/inward/record.url?eid=2-s2.0-0031714655&partnerID=tZOtx3y1

Pederick, L. A. (1968). Chromosome inversions in Pinus radiata. Silvae Genet, 17, 22–26.

Pritchard, J. K., Stephens, M., & Donnelly, P. (2000). Inference of population structure using multilocus genotype data. Genetics, 155(2), 945–959.

Puig, M., Casillas, S., Villatoro, S., & Cáceres, M. (2015). Human inversions and their functional consequences. Briefings in Functional Genomics, 14(5), 369–379. doi:10.1093/bfgp/elv020

Pyhäjärvi, T, Garcia-Gil, M. R., Knurr, T., Mikkonen, M., Wachowiak, W., & Savolainen, O. (2007). Demographic history has influenced nucleotide diversity in European *Pinus sylvestris* populations. Genetics, 177(3), 1713–1724. doi:10.1534/genetics.107.077099

Pyhäjärvi, Tanja, García-Gil, M. R., Knürr, T., Mikkonen, M., Wachowiak, W., & Savolainen, O. (2007). Demographic history has influenced nucleotide diversity in European Pinus sylvestris populations. Genetics, 177(3), 1713–1724. doi:10.1534/genetics.107.077099

Pyhäjärvi, Tanja, Hufford, M. B., Mezmouk, S., & Ross-Ibarra, J. (2013). Complex patterns of local adaptation in teosinte. Genome Biology and Evolution, 5(9), 1594–1609. doi:10.1093/gbe/evt109

Racimo, F., Berg, J. J., & Pickrell, J. K. (2018). Detecting polygenic adaptation in admixture graphs. Genetics, 208(4), 1565–1584.

Rockman, M. V. (2012). The QTN program and the alleles that matter for evolution: All that’s gold does not glitter. Evolution, 66(1), 1–17. doi:10.1111/j.1558-5646.2011.01486.x

Savolainen, O., Lascoux, M., & Merilä, J. (2013). Ecological genomics of local adaptation. Nature Reviews. Genetics, 14(11), 807–20. doi:10.1038/nrg3522

Savolainen, O., Pyhäjärvi, T., & Knürr, T. (2007). Gene Flow and Local Adaptation in Trees. Annual Review of Ecology, Evolution, and Systematics, 38(1), 595–619. doi:10.1146/annurev.ecolsys.38.091206.095646

Schmidt, P. S., Zhu, C.-T., Das, J., Batavia, M., Yang, L., & Eanes, W. F. (2008). An amino acid polymorphism in the couch potato gene forms the basis for climatic adaptation in Drosophila melanogaster. Proceedings of the National Academy of Sciences, 105(42), 16207–16211.

Slatkin, M. (1973). GENE FLOW AND SELECTION IN A CLINE. Genetics, 75(4), 733–756.

Sundell, D., Mannapperuma, C., Netotea, S., Delhomme, N., Lin, Y., Sj, A., … Street, N. R. (2015). The Plant Genome Integrative Explorer Resource: PlantGenIE . org, 1149–1156.

Tajima, F. (1989). Statistical method for testing the neutral mutation hypothesis by DNA polymorphism. Genetics, 123(3), 585–595. doi:PMC1203831

Thompson, M. J., & Jiggins, C. D. (2014). Supergenes and their role in evolution. Heredity, (August 2013), 1–8. doi:10.1038/hdy.2014.20

Thorvaldsdóttir, H., Robinson, J. T., & Mesirov, J. P. (2013). Integrative Genomics Viewer (IGV): High-performance genomics data visualization and exploration. Briefings in Bioinformatics, 14(2), 178–192. doi:10.1093/bib/bbs017

Tyrmi, J. S. (2018). STAPLER: a simple tool for creating, managing and parallelizing common high-throughput sequencing workflows. BioRxiv, 445056. doi:10.1101/445056

Vitalis, R., Gautier, M., Dawson, K. J., & Beaumont, M. a. (2014). Detecting and measuring selection from gene frequency data. Genetics, 196(3), 799–817. doi:10.1534/genetics.113.152991

Wachowiak, W., Balk, P. A., & Savolainen, O. (2009). Search for nucleotide diversity patterns of local adaptation in dehydrins and other cold-related candidate genes in Scots pine (Pinus sylvestris L.). Tree Genetics & Genomes, 5(1), 117–132. doi:10.1007/s11295-008-0188-3

Wellenreuther, M., & Bernatchez, L. (2018). Eco-Evolutionary Genomics of Chromosomal Inversions. Trends in Ecology and Evolution, 33(6), 427–440. doi:10.1016/j.tree.2018.04.002

Westbrook, J. W., Neves, L. G., Kirst, M., Peter, G. F., Chamala, S., Nelson, C. D., … Echt, C. S. (2015). A Consensus Genetic Map for Pinus taeda and Pinus elliottii and Extent of Linkage Disequilibrium in Two Genotype-Phenotype Discovery Populations of Pinus taeda. G3: Genes|Genomes|Genetics, 5(8), 1685–1694. doi:10.1534/g3.115.019588

Wu, T. D., & Nacu, S. (2010). Fast and SNP-tolerant detection of complex variants and splicing in short reads. Bioinformatics, 26(7), 873–881.

Yeaman, S. (2013). Genomic rearrangements and the evolution of clusters of locally adaptive loci. Proceedings of the National Academy of Sciences, 110(19), E1743–E1751. doi:10.1073/pnas.1219381110

Yeaman, Sam. (2015). Local Adaptation by Alleles of Small Effect. The American Naturalist, 186(S1), S74–S89. doi:10.1086/682405

Yeaman, Sam, & Whitlock, M. C. (2011). The genetic architecture of adaptation under migration-selection balance. Evolution; International Journal of Organic Evolution, 65(7), 1897–911. doi:10.1111/j.1558-5646.2011.01269.x

Zonneveld, B. J. M. (2012). Conifer genome sizes of 172 species, covering 64 of 67 genera, range from 8 to 72 picogram. Nordic Journal of Botany, 30(4), 490–502. doi:10.1111/j.1756-1051.2012.01516.x

