## Supplement for "Genomics of clinal local adaptation in *Pinus sylvestris* under continuous environmental and spatial genetic setting"

### Table of Contents:


| **Supplementary methods** | Page 2 |
|---|---|
| **Table S1** | Page 4 |
| **Table S2** | Page 5 |
| **Figure S1** | Page 6 |
| **Figure S2** | Page 7 |
| **Figure S3** | Page 8 |
| **Figure S4** | Page 9 |


### Supplementary methods

Alignment and SNP calling are often challenging in conifers as due to the repetitive nature of their genomes (Wegrzyn et al., 2014), leading to errors in read alignment. Alignments of individual samples visualized with IGV (Thorvaldsdottir, James, & Jill, 2012) revealed that large proportion of target area contained not only reads with probably correct alignment (few or no mismatches to reference genome) but in addition reads which were likely incorrectly aligned to the area (multiple mismatches and gaps). This suggests that the off-target, paralogous sequence had been captured along with the target sequence.

Areas containing paralogous sequence may lead to spurious SNP calls inflating the number of SNPs and distorting the shape of the allele frequency spectrum. Identification of such areas is possible since overlap of correctly and incorrectly aligned reads may be detected as heterozygous genotype calls, which are not expected with haploid DNA that was used for sequencing. To alleviate the incorrect alignment issue the SNP calling step was performed twice. The first SNP call to detect spurious heterozygous areas was performed with freebayes (Garrison & Marth, 2012) using parameter --ploidy 2 to detect heterozygous variant calls. Other parameters were set as -T 0.01 --min-coverage 5 -u -X. Areas containing heterozygous genotypes and nearby areas with radius of 25 bases were then marked to a bed file for later filtering as paralogous areas may also contain spurious homozygous SNP calls which should be removed. Visualizations of alignments indicated that paralogous areas were often saturated with heterozygous SNP calls hence a filter radius of 25 bases was deemed sufficiently large to filter out possible spurious homozygous calls. A bed file defining non-paralogous areas was then created by generating a bed file for the whole *P. taeda* reference genome and removing the paralogous areas with bedtools (Quinlan & Hall, 2010) subtract command.

Variant calling was redone to achieve the final set of SNPs with the same parameters as in the first call, but --ploidy 1 argument was used instead and the bed file defining non-paralogous areas was provided as target area with -t parameter. As some downstream analysis requires information on the length of the sequence available for analysis (henceforth referenced as 'available genome') a VCF file containing genotype calls including monomorphic sites was generated by including --report-monomorphic parameter to freebayes command line. As calling monomorphic genotypes for large number of samples requires excessive amount of RAM and a long runtime, the SNP calling was performed only for scaffolds longer than 1000 bases. This removes only 14.3 % of total sequence but the total number of scaffolds is reduced from over 14 million to one million greatly reducing the computational runtime of downstream analysis. No exome capture baits had been designed for the omitted areas. The genotype calling was parallelized by performing the run separately for each sample in its own thread. The produced VCF files were then combined with vcflib vcfcombine tool.

| Gene name | Scaffold name | Position in scaffold | Description |
|---|---|---|---|
| LURP1 | C32491382 | 40265, 40280 | Associated to parasite defence in A. thaliana (Knoth & Eulgem, 2008) |
| Auxin response factor-5 like | C32502832 | 26212 | Involved in salt and drought tolerance in A. thaliana (Kang et al., 2018), regulates fruit set and development in tomato (Liu et al., 2018) |
| Tetratricopeptide repeat domain (TPR) | scaffold291986.2 | 49639, 49782, 50073, 50079 | Possibly associated to osmotic stress responses (Rosado et al., 2006), and shown to be outlier in environmental association tests in Eucalyptus microcarpa (Jordan, Hoffmann, Dillon, & Prober, 2017) |
| Gamma carbonic anhydrase 1 | scaffold204210 | 124240, 124265, 124403 | Carbonic anhydrases have various roles in photosynthesis where they work in conjunction with Rubisco (Badger & Price, 1994) |
| 51 SNPs within Pentatricopeptide repeat containing genes | C32563588<br>C32567682<br>scaffold29782<br>scaffold29782<br>scaffold39072<br>scaffold160879<br><br>scaffold274519<br>scaffold535265<br><br>scaffold537986<br>scaffold565735<br>scaffold876405<br>scaffold901805<br>tscaffold3453<br>tscaffold4081<br>tscaffold4747<br>tscaffold6603<br>tscaffold7974<br>tscaffold8936 | 99631<br>162184<br>20068<br>20192<br>120552<br>101352, 101517, 101607, 102048, 102086, 102159,<br>102600, 102615, 102671, 102788, 102855, 103194,<br>103331, 103332<br>55201, 55213<br>41272, 41380, 41440, 41636, 41746, 41785, 41893, 41952,<br>42298, 42300, 42334, 42363, 42523, 42535<br>52711, 52942<br>49168<br>27038<br>73961<br>149057<br>216057<br>539514<br>41052, 113519<br>403268, 403301, 403357, 403453, 403468<br>213987 | They have multitude of tasks in plants (Barkan & Small, 2014) and have been shown to be outliers in selection scans e.g. in A. lyrata (Foxe & Wright, 2009) and in conifers (Scalfi et al., 2014; Yeaman et al., 2016) |
| G−type lectin S−receptor−like serine/threonine−protein kinase | scaffold310452 | 41495 | May have a role in response to salt and drought stress (Sun et al., 2013) |
| myb family transcription factor APL isoform X2 | scaffold310452 | 41495 | Various roles in biotic and abiotic stresses (Ambawat, Sharma, Yadav, & Yadav, 2013) |
| erd1 | tscaffold1451 | 366226, 366447 | Associated with dehydration stress (Simpson et al., 2003) |

*Table S1. Examples of interesting outliers detected in the linear regression outlier analysis. References mentioned in the Description column are listed below.*

| Population | Inari | Kolari | Kalvia | Punkaharju | Kalsnava | Radom | Ust-Chilma | Megdurechensk | Ust-Kulom | Penzenskaja | Volgogradskaja |
|---|---|---|---|---|---|---|---|---|---|---|---|
| **Kolari** | 0.014 | | | | | | | | | | |
| **Kalvia** | 0.000 | 0.016 | | | | | | | | | |
| **Punkaharju** | 0.000 | 0.017 | 0.005 | | | | | | | | |
| **Kalsnava** | 0.019 | 0.019 | 0.023 | 0.020 | | | | | | | |
| **Radom** | 0.002 | 0.020 | 0.005 | 0.007 | 0.015 | | | | | | |
| **Ust-Chilma** | 0.010 | 0.022 | 0.013 | 0.010 | 0.034 | 0.018 | | | | | |
| **Megdurechensk** | 0.008 | 0.020 | 0.011 | 0.009 | 0.029 | 0.016 | 0.001 | | | | |
| **Ust-Kulom** | 0.056 | 0.040 | 0.056 | 0.057 | 0.041 | 0.045 | 0.050 | 0.048 | | | |
| **Penzenskaja** | 0.008 | 0.023 | 0.011 | 0.013 | 0.033 | 0.015 | 0.009 | 0.009 | 0.056 | | |
| **Volgogradskaja** | 0.000 | 0.017 | 0.004 | 0.005 | 0.022 | 0.007 | 0.010 | 0.006 | 0.056 | 0.001 | |
| **Baza** | 0.068 | 0.081 | 0.073 | 0.071 | 0.077 | 0.065 | 0.083 | 0.077 | 0.113 | 0.081 | 0.072 |

*Table S2. Weighted genome-wide averages of pairwise $F_{ST}$ estimates for all populations but with genomic areas identified as part of the haplotype structure omitted.*

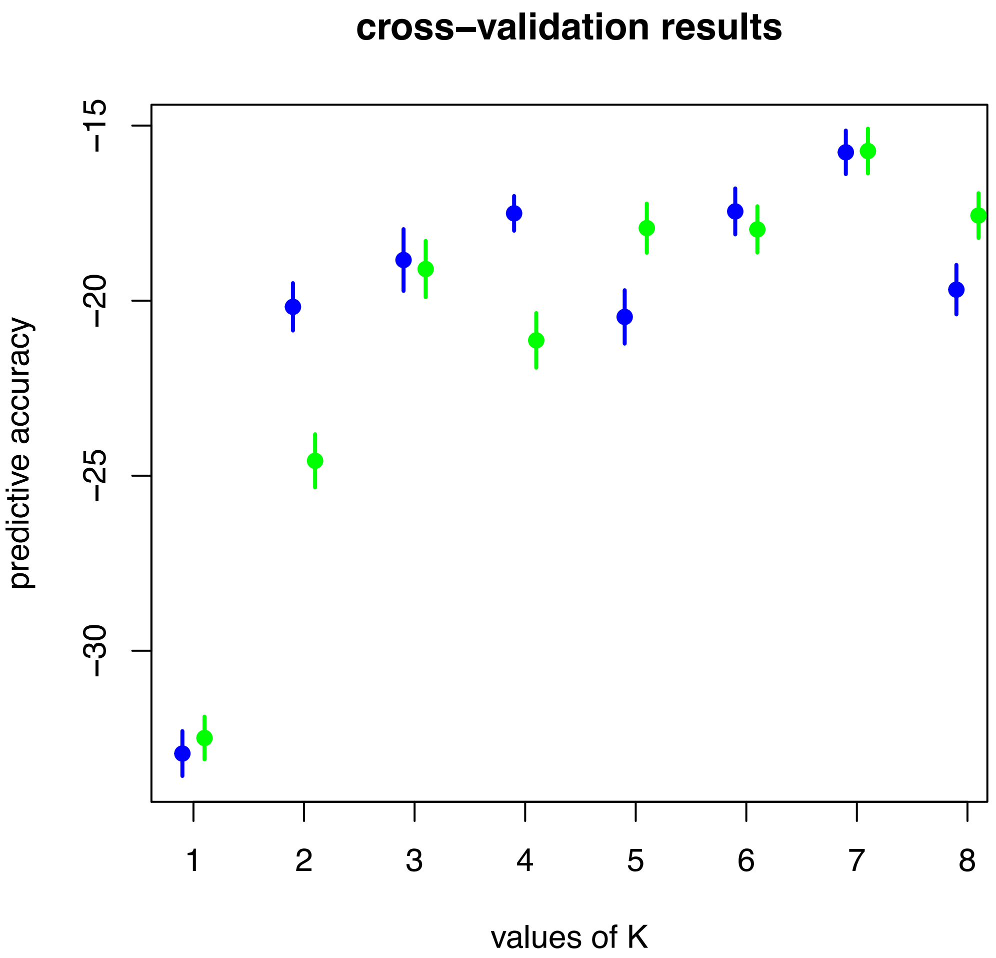


*Figure S1. Cross-validation results for conStruct for K-values 1 to 8 for spatial (blue) and non-spatial (green) models using 40 iterations.*

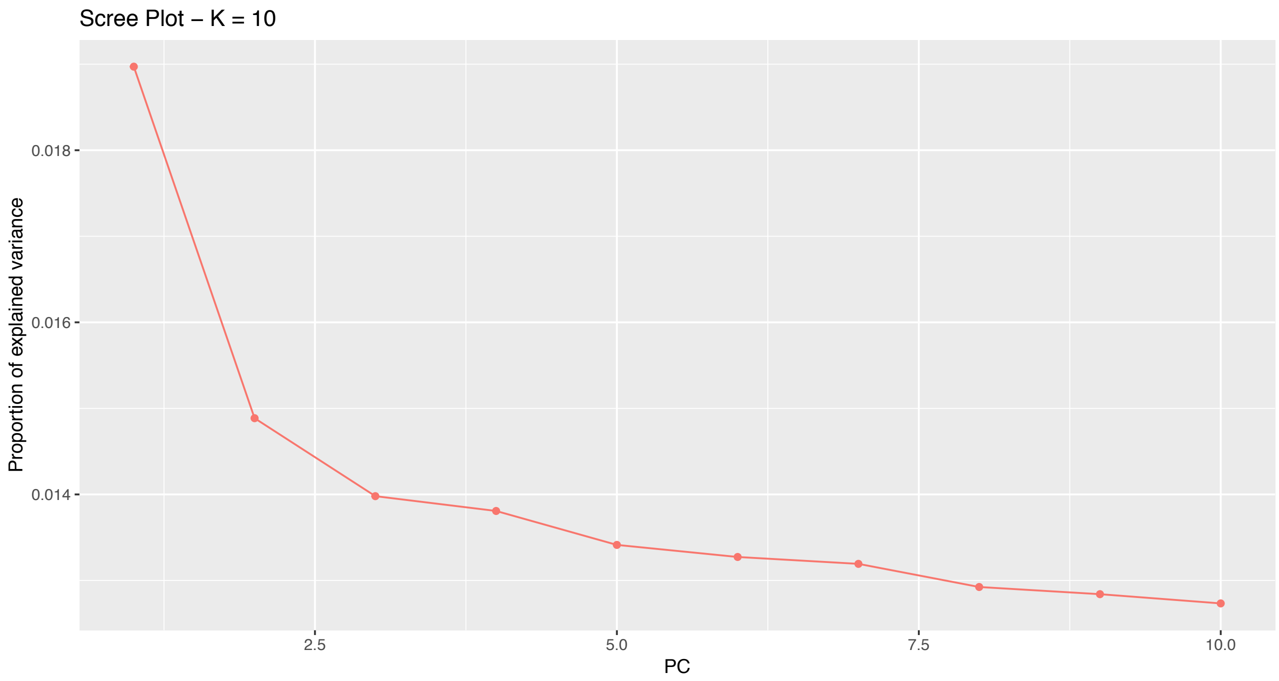


*Figure S2. Screeplot procuded with pcadapt. Here the eigenvalues corresponding to random variation lie on the straight line and the values corresponding to population structure depart from it.*

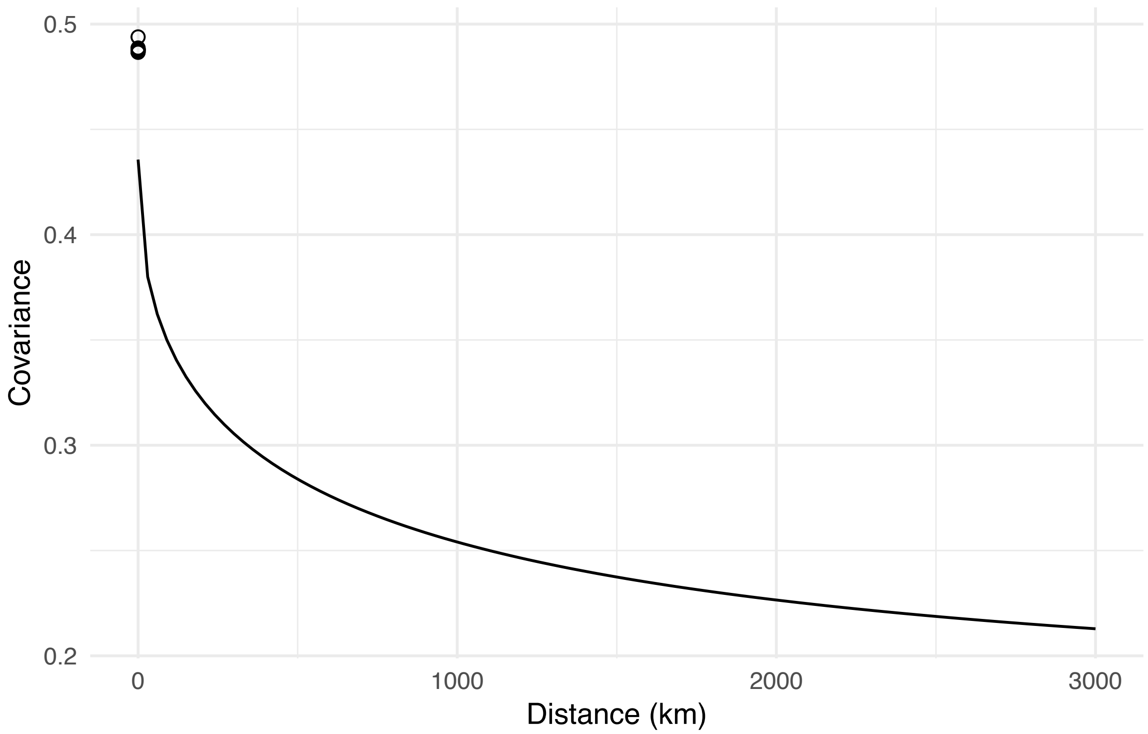


*Figure S3. The within population covariance (dots) and the decrease of covariance between populations as between-population distance increases (line) for the first layer for the conStruct analysis.*

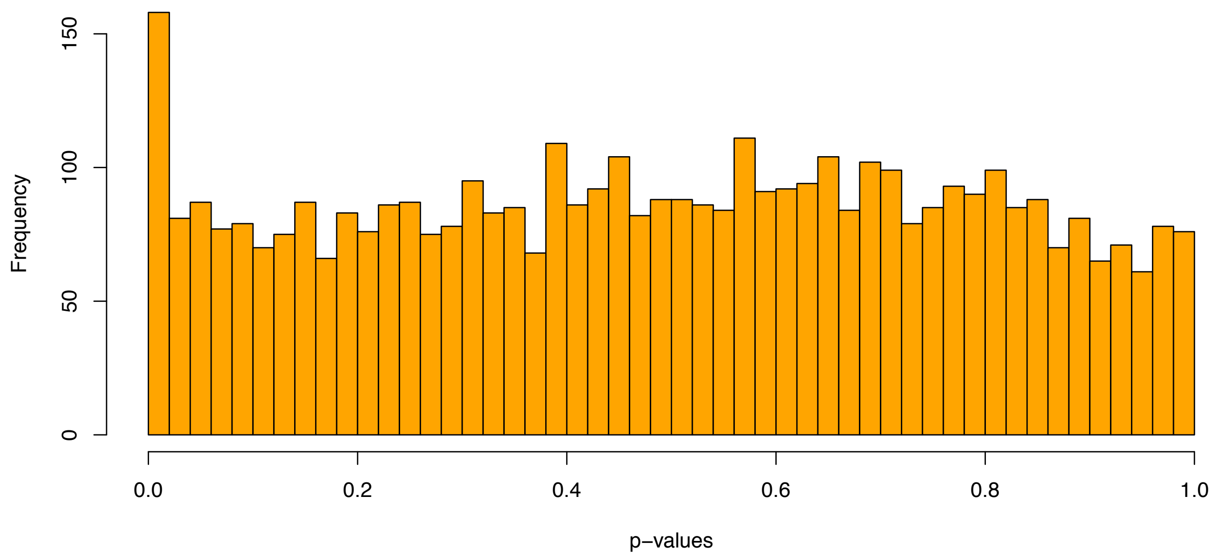


*Figure S4. Distribution of p-values for pcadapt analysis. Excess of low p-values indicates the presence of outliers.*
